# Antagonistic conflict between transposon-encoded introns and guide RNAs

**DOI:** 10.1101/2023.11.20.567912

**Authors:** Rimantė Žedaveinytė, Chance Meers, Hoang C. Le, Edan E. Mortman, Stephen Tang, George D. Lampe, Sanjana R. Pesari, Diego R. Gelsinger, Tanner Wiegand, Samuel H. Sternberg

**Affiliations:** Department of Biochemistry and Molecular Biophysics, Columbia University; New York, NY 10032, USA; Department of Genetics and Development, Columbia University; New York, NY 10032, USA; Biochemistry and Molecular Biophysics Program, University of California, San Diego, CA, USA

## Abstract

TnpB nucleases represent the evolutionary precursors to CRISPR-Cas12 and are widespread in all domains of life, presumably due to the critical roles they play in transposon proliferation. IS605-family TnpB homologs function in bacteria as programmable homing endonucleases by exploiting transposon-encoded guide RNAs to cleave vacant genomic sites, thereby driving transposon maintenance through DSB-stimulated homologous recombination. Whether this pathway is conserved in other genetic contexts, and in association with other transposases, is unknown. Here we uncover molecular mechanisms of transposition and RNA-guided DNA cleavage by IS607-family elements that, remarkably, also encode catalytic, self-splicing group I introns. After reconstituting and systematically investigating each of these biochemical activities for a candidate ‘IStron’ derived from *Clostridium botulinum*, we discovered sequence and structural features of the transposon-encoded RNA that satisfy molecular requirements of a group I intron and TnpB guide RNA, while still retaining the ability to be faithfully mobilized at the DNA level by the TnpA transposase. Strikingly, intron splicing was strongly repressed not only by TnpB, but also by the secondary structure of ωRNA alone, allowing the element to carefully control the relative levels of spliced products versus functional guide RNAs. Our results suggest that IStron transcripts have evolved a sensitive equilibrium to balance competing and mutually exclusive activities that promote transposon maintenance while limiting adverse fitness costs on the host. Collectively, this work explains how diverse enzymatic activities emerged during the selfish spread of IS607-family elements and highlights molecular innovation in the multi-functional utility of transposon-encoded noncoding RNAs.

## INTRODUCTION

Transposases are among the most abundant genes in nature, due to their high copy number and cross-kingdom host diversity (*1*). These enzymes transpose DNA through chemically disparate mechanisms and can be classified based on the presence of conserved domains that fall into one of several major protein families, including DD(E/D), tyrosine transposase, and serine transposase (*2*, *3*). In turn, the DNA elements recognized by transposases bear hallmark sequence features at their left and right ends, ensuring a tight molecular coupling between transposon substrates and the enzymes that mobilize them (*4*). Interestingly, two large bacterial transposon families typified by IS605 and IS607 encode distinct transposase machineries, but the very same accessory factor, TnpB (**Fig. 1A**) (*3*, *5–9*). Eukaryotic TnpB homologs, known as Fanzors, are associated with an even greater diversity of transposons/transposases, highlighting the pervasive and recurrent co-option events involving this ubiquitous gene, as well as the useful properties it apparently provided to these mobile elements over evolutionary timescales (*10–13*). TnpB co-option and host domestication also notably led to the evolution of type II and type V CRISPR-Cas adaptive immunity, as both Cas9 and Cas12 RNA-guided nucleases can be phylogenetically linked to these transposon-encoded accessory factors (*5*, *6*, *8*, *9*).

**Fig. 1.**
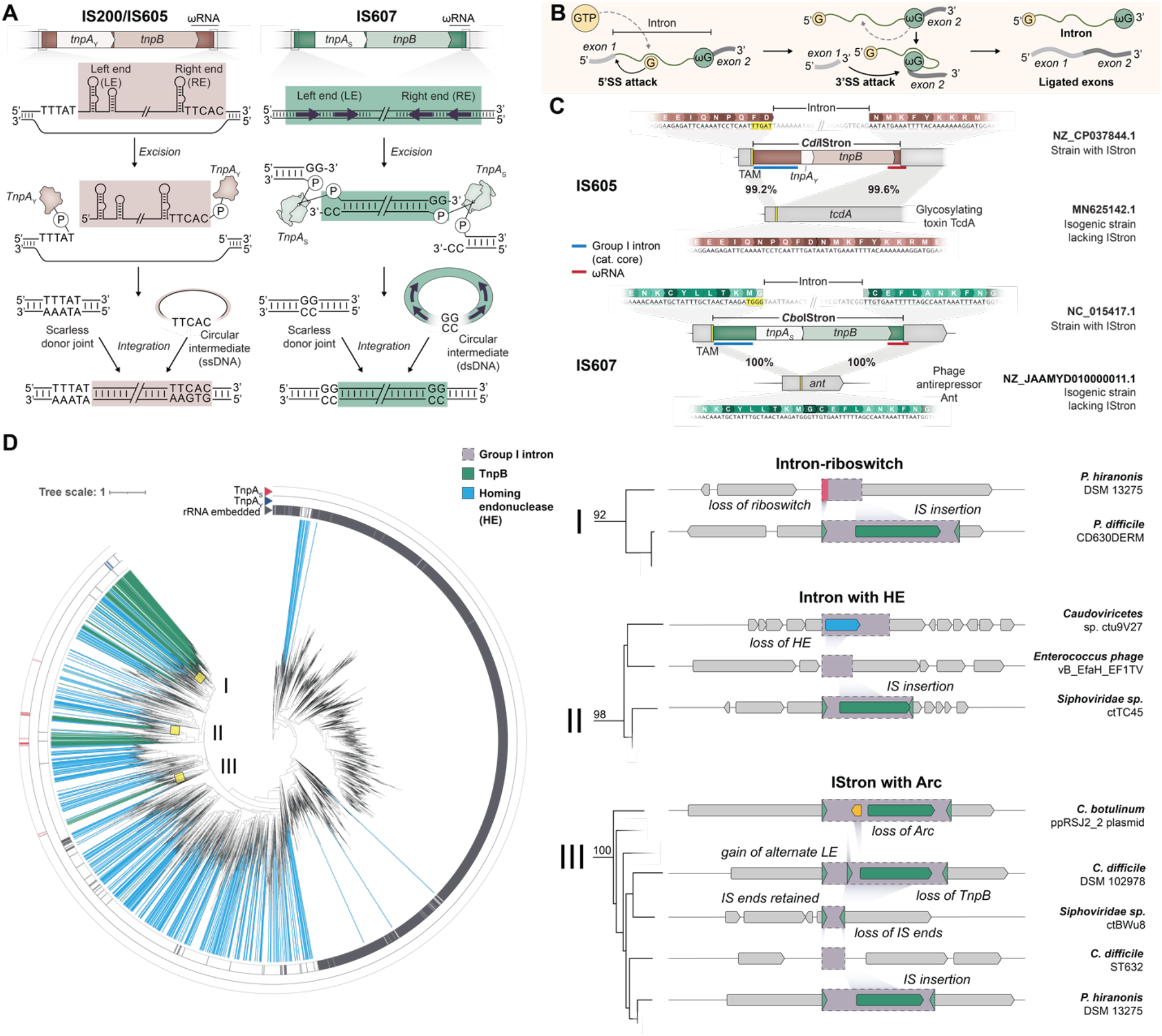
Genomic architecture and endogenous splicing activity of TnpB-encoding IStrons. (**A**) IS200/IS605-family transposons mobilize through a ssDNA intermediate using a tyrosine-family recombinase (TnpA_Y_, left); in contrast, IS607-family transposons mobilize through a dsDNA intermediate using a serine-family transposase (TnpA_S_, right). Transposons of both families are bounded by conserved left end (LE) and right end (RE) sequences, encode *tnpB* accessory genes, excise as circular intermediates, and generate scarless donor joints that precisely regenerate the native genomic sequence. (**B**) Schematic of the group I intron self-splicing pathway, in which binding of exogenous GTP leads to consecutive transesterification reactions to yield ligated exons and the excised intron RNA. Introns often encode their own ORFs. (**C**) Genetic architecture of representative IS605 and IS607-family IStrons from *Clostridia* strains (top), alongside homologous sites from related strains that lack the transposon insertion (bottom). The percent sequence identity between shaded regions (grey) is shown, as are the genomic accession IDs; TAMs are highlighted in yellow. Flanking exons are labeled, and the inset (middle) highlights the predicted mRNA splicing products that regenerate a functional open reading frame (ORF). (D) Phylogenetic tree of group I introns that are structurally related to the *Cbo*IStron group I intron (left), with genetic architectures of selected clades schematized (right). The outer rings of the tree indicate associations with TnpA_S_ or TnpA_Y_, as well as whether the group I intron is encoded within an rRNA locus. The green and blue colors indicate associations with TnpB nucleases or homing endonucleases (HE), which fall into many distinct enzyme families (LAGLIDADG, GIY-YIG, HNH, His-Cys Box, and Endonuclease VII). Bootstrap values are indicated for select nodes.

We recently showed that TnpB and guide RNAs play a key role in the maintenance and spread of IS200/IS605-family transposons through a peel-and-paste, cut-and-copy transposition mechanism that involves RNA-guided DNA cleavage (*7*). A key insight resulted from our observations that the TnpA — tyrosine-family recombinase responsible for IS200/IS605 mobilization — (hereafter TnpA_Y_) catalyzes a higher frequency of transposon excision than integration, and that scarless excision leads to permanent loss of the element at the donor site — foreboding eventual extinction of the mobile element. In the presence of TnpB, though, targeted DNA double-strand breaks (DSBs) trigger efficient recombination with a sister chromosome still harboring the transposon, thereby promoting retention. We demonstrated this biological function for representative IS605 elements from *Geobacillus stearothermophilus*, but intriguingly, our phylogenetic analyses uncovered the presence of homologous TnpB genes in another family of transposons, known as IS607 (*7*).

IS607 transposons are among the most widespread transposable elements in nature, spanning bacteria, archaea, eukaryotes, and are even found in viruses that infect these hosts (*14–16*). They are mobilized by a serine-family transposase also named TnpA (hereafter TnpA_S_) via a recombination reaction akin to small and large serine recombinases like Tn3 and Bxb1, respectively (*17–19*). Despite the common naming scheme, TnpA_Y_ and TnpA_S_ do not share a common ancestor and employ distinct catalytic mechanisms and substrate preferences: whereas TnpA_Y_ transposes IS200/IS605-family elements through a circular single-stranded DNA (ssDNA) intermediate using a 3’-phospho-tyrosine linkage (*20*, *21*), TnpA_S_ transposes IS607-family elements through a circular double-stranded DNA (dsDNA) intermediate using a 5’-phospho-serine linkage (*4*, *16*, *22*) (**Fig. 1A**). Yet a common feature of both pathways is the role that the transposase plays in chemically resealing the sequences flanking the transposon at the donor site during the excision step, to produce a scarless ‘donor joint’ that regenerates the original genomic sequence (**Fig. 1A**). Thus, IS607-family transposons (hereafter simply IS607), like IS200/IS605-family transposons (hereafter simply IS605), are at risk of copy number loss and eventual extinction, under circumstances where the excision frequency outpaces the integration frequency at new target sites (*7*). We therefore hypothesized that TnpB might satisfy the very same evolutionary purpose for both types of elements, to promote their maintenance via a programmable cut-and-copy mechanism.

A further molecular feature caught our attention: the presence of conspicuous group I introns inside a subset of both IS605 and IS607 transposons (**Fig. 1B,C**). Introns are often themselves considered mobile genetic elements, and they possess the exceptional ability to persist silently in genomes by removing themselves from interrupted transcripts, thereby restoring contiguous structural RNA or protein-coding genes (*23*). Group I introns are large, ancient RNA enzymes, or ribozymes, that require only Mg^2+^ and GTP for self-splicing (*24*) (**Fig. 1B**), and unlike group II introns, which utilize protein cofactors (maturases) for splicing and reverse transcriptases for mobilization (*25*), some group I introns are thought to use reverse splicing for mobilization (*26*, *27*). A notable exception are introns that encode homing endonucleases, which promote intron mobility through DNA cleavage and recombination at fixed target sites (*28–30*). Based on the genetic architecture of group I introns that also encode TnpA and TnpB, and on the presence of predicted transposon end sequences (**Fig. 1C**), we hypothesized that these elements – termed IStrons (*31–35*) – would have the ability to mobilize via DNA transposition while mitigating fitness costs via splicing. Intriguingly, though, the TnpB-associated guide RNAs encoded by IStrons, termed ωRNAs (*8*, *9*), would presumably span the intron-exon junction, and thus compete directly with the process of splicing itself. We wondered how multi-functional IStron transcripts can fulfill the molecular requirements for both RNA splicing and RNA-guided DNA targeting, while also retaining DNA sequences that are productively transposed by TnpA.

Focusing on IS607-family elements for which the role of TnpB has not been explored, here we reconstituted DNA transposition, RNA self-splicing, and RNA-guided DNA cleavage activities of a representative IStron from *Clostridium botulinum*, the causative agent of botulism (*36–38*). Our experiments revealed the overlapping and contingent molecular determinants of three distinct nucleic acid scission/joining reactions, highlighting the remarkable ability for a single polynucleotide sequence to direct orthogonal and multifunctional chemical reactions. In particular, we found that the full-length IStron transcript serves parallel but mutually exclusive roles as an intron capable of protein-independent self-splicing, and as a non-coding guide RNA that specifies genomic target sites for DNA cleavage. The relevant balance between these activities is controlled both by intrinsic structural features of the RNA itself, and by the TnpB nuclease, highlighting its central regulatory role in the IStron lifecycle. Collectively, our work unveils an ingenious and unprecedented convergence of molecular activities that exemplify the capacity for molecular innovation by transposable elements.

## RESULTS

### Bioinformatics analyses of IStron elements

IStrons feature unusually long intergenic regions upstream of *tnpA*, which are clearly contained within the transposon boundaries and encode a structured ncRNA that matches the self-splicing group I intron covariance model from the Rfam database (*39*) (**Fig. 1C**). When we systematically analyzed the genomic neighborhood of all *tnpB* genes, we identified numerous clades that were intron-associated, with clear evidence of independent ancestral co-option events between catalytic ribozymes and distinct IS605 (*tnpA*_Y_) or IS607 (*tnpA*_S_) transposons (**fig. S1**). Motivated by this observation, we constructed an improved group I intron covariance model (CM), based on specific IStron-encoded group I introns, and used it to identify all related group I introns in the NT database. This analysis highlighted diverse genomic contexts and associations with an eclectic mix of auxiliary genes, including TnpA transposases, TnpB nucleases, homing endonucleases, and uncharacterized accessory factors (**Fig. 1D**). Closer examination illuminated dynamic and recurring gene gain and loss events between otherwise closely related introns, highlighting the rapid diversification of these elements. We also uncovered a novel IStron-encoded ORF with a predicted homodimeric ribbon-helix-helix (RHH2) structure reminiscent of Arc-family transcription factors (*40*) (**Fig. 1D, fig. S2A**). A broader phylogenetic analysis revealed hundreds of additional Arc-like homologs associated not only with TnpB, but also with homing endonucleases, other mobile genetic elements, and various toxin-antitoxin systems (**fig. S2B**). Previously studied Arc homologs, including ω (*41*), CopG (*42*), and MetJ (*43*, *44*), have been shown to recognize arrays of direct or inverted repeat sequences, suggesting a potential role in regulating transposition, gene transcription, or RNA biogenesis, possibly in conjunction with TnpA, TnpB, or the intron.

We focused our subsequent analyses on representative IS605- and IS607-family IStrons from two biomedically relevant strains of Clostridia, *Clostridioides difficile* (*Cdi*) and *Clostridium botulinum* (*Cbo*), respectively. *C. difficile* strain 630 harbors 8 identifiable IStron copies within the IS605 family (hereafter *Cdi*IStrons), alongside three freestanding group I introns and three intron-less IS605 elements (**fig. S3A, fig. S4A, and table S1**). None of the *Cdi*IStrons encode full-length TnpA, indicating that these elements are non-autonomous or immobile, but the presence of *tnpA*_Y_ gene fragments allowed us to confidently place them within the IS605 superfamily. We were able to easily identify transposon left end (LE) and right end (RE) boundaries using CMs and comparative genomics (**Fig. 1C and fig. S3B,C**). These boundary assignments were further corroborated by the presence of a conserved 5′-TTGAT-3′ transposon-adjacent motif (TAM) abutting the LE (**fig. S3D**), a sequence similar to the TAM we previously defined for related TnpA_Y_ and TnpB homologs (*7*). The group I intron falls within subtype A2 (*31*, *34*, *45–47*) and exhibits predicted secondary structure features that match with the previously characterized Twort ribozyme (*48*, *49*) (**fig. S3E**). Putative 5′ and 3′ splice sites (5′SS and 3′SS) were identified based on their anticipated coincidence with the transposon LE/RE boundaries, and by closer inspection of the genes interrupted by the IStron element, since the anticipated exon-exon splicing product perfectly regenerated the wild-type gene coding sequence (**Fig. 1C**). Finally, we identified transposon-encoded ωRNAs using a new *Cdi*IStron-anchored CM (**fig. S5E**) and found that these ωRNAs were consistently located in the 3′-UTR of *tnpB*, similarly to IS605-family transposons lacking group I introns (**Fig. 1C and fig. S3F**). Importantly, ωRNAs presumably encompass the same regions that are critical for 3′SS recognition by the group I intron, suggesting potential competition between self-splicing and guide RNA binding.

We mined published *C. difficile* RNA-sequencing data (*50*, *51*) and observed expression from all intron and IS605 elements, though precise mapping to predicted ωRNA regions was confounded by their highly repetitive nature and the sequencing method (**fig. S4**). The dual presence of unspliced and spliced reads for all but one of the *Cdi*IStron copies provided concrete evidence of intron splicing, with exon-exon junctions that perfectly matched our annotated transposon LE/RE boundaries (**Fig. 1C and fig. S4B**). Quantification of splicing activity revealed a surprisingly wide range of efficiencies, despite extremely high group I intron sequence conservation across all 8 IStron copies (**fig. S4A,B**), suggesting that genomic context may play an important role in regulating intron folding, intron catalysis, and/or RNA stability.

Turning our attention to IS607-family elements, for which TnpB and transposition have not been extensively studied, we identified an IStron in *C. botulinum* strain BKT015925 (hereafter *Cbo*IStron) that encoded a full-length TnpB and a serine recombinase (TnpA_S_). Intriguingly, the *Cbo*IStron is located on a large extrachromosomal plasmid, alongside an IStron lacking TnpA, multiple additional IS607 elements, and the botulinum neurotoxin (**fig. S5A and table S1**) (*37*, *38*). IS607 transposons lack inverted repeats or imperfect palindromic ends, and transposition does not generate a target-site duplication (TSD) (*16*, *22*, *52*), rendering unbiased transposon boundary detection with sequence/structure-based models challenging. However, we were able to provisionally assign LE/RE boundaries for these elements using comparative genomics (**Fig. 1C and fig. S5B**), which, together with the analysis of homologous *Cbo*IStrons, revealed a predicted guanine-rich TAM motif harboring a consensus sequence of 5′-TGGG-3′ (**fig. S5C**). *C. botulinum* group I introns are similar to *C. difficile* group I introns in their predicted structure (**fig. S5D**), and indeed could be detected using the same CM. The putative splice sites again overlapped with the transposon DNA boundaries, based on analysis of the flanking exonic gene fragments (**Fig. 1C**).

We were especially interested to inspect the putative *Cbo*IStron ωRNA. In the case of IS605 transposons, conserved ωRNA stem-loops bound by TnpB coincide with palindromic DNA stem-loops in the RE (**fig. S3F,G**), which are specifically recognized by TnpA_Y_ during transposon excision and integration (**Fig. 1A**) (*53*). IS607 transposons, on the other hand, are recognized as dsDNA by TnpA_S_ through poorly understood sequence determinants (*16*, *22*, *52*), suggesting an alternative evolution of RE sequences that would satisfy requirements of both TnpA_S_ (DNA) and TnpB (RNA). After identifying the likely ωRNA through a *Cbo*TnpB-anchored CM (**fig. S5E**), we found that, remarkably, ωRNAs exhibit universal structural features despite their association with distinct IS605 and IS607 elements, characterized by three consecutive stem-loops (SL) upstream of the guide sequence that comprise the ωRNA ‘scaffold’ (**fig. S5F**). The first SL, termed the nexus, is predicted to base-pair with complementary sequences at the scaffold 3′ end via a pseudoknot (PK) interaction (**fig. S5F**), and this feature is conserved across characterized TnpB-ωRNA systems, as well as the guide RNAs for Cas12-family nucleases (*54–58*). These commonalities attest to the evolutionary relatedness of TnpB and Cas12 proteins and their RNA substrates, and demonstrate that essential RNA structural motifs can be accommodated by unique DNA requirements across diverse transposable elements.

We next set out to reconstitute the enzymatic activities of TnpA_S_, TnpB, and a group I intron, as a way to probe the overlapping and potentially antagonistic processes of RNA-guided DNA cleavage and RNA splicing. We decided to focus exclusively on the IS607-family *Cbo*IStron, in order to also determine whether TnpB plays a similar role in transposon maintenance for this ubiquitous family of elements, as compared to IS200/IS605-family transposons.

### TnpA_S_ catalyzes efficient IStron excision and integration

We started by developing a heterologous transposon excision assay in *E. coli*, in which cells were transformed with a *Cbo*TnpA_S_ expression plasmid (pTnpA_S_) and a transposon donor plasmid (pDonor) encoding a minimal *Cbo*IStron (mini-IS) lacking *tnpA*_S_ and *tnpB* genes. Transposon excision is expected to generate a circularized dsDNA transposon intermediate and regenerated donor site containing the adjoined TAM and target site (donor joint), which we detected using either endpoint or quantitative PCR (**Fig. 2A and fig. S6A**). TnpA_S_ catalyzed robust transposon excision, an activity that was ablated in the presence of protein mutations to the TnpA_S_ active site, DNA mutations to mismatch the core dinucleotides or remove the transposon ends (**Fig. 2B**). Sanger sequencing revealed that plasmid excision products featured precise donor joints (**Fig. 2C**), and we also readily detected the presence of circular excision intermediates (**fig. S6B**). DNA excision efficiencies were ∼5% after culturing overnight (**Fig. 2D**), and thus orders of magnitude higher than TnpA_Y_ from IS605 elements (*7*), suggesting that excisional transposon loss might be an even greater risk for IS607 elements. To more accurately define the minimum necessary transposon end sequences required for TnpA_S_-DNA recognition, we quantified the effects of serial LE/RE deletions on excision. These experiments revealed that only 40- and 60-bp of the LE and RE were critical, respectively (**Fig. 2D and fig. S6C**). Next, we analyzed sequence conservation within both transposon ends and identified a series of short, 6-7-bp inverted motifs that we hypothesized would be important for TnpA_S_ binding. Mutating any one of these motifs alone had variable effects, but concurrent disruption to at least two motifs completely abolished excision activity (**fig. S6D**), supporting their importance for proper recognition and assembly of an enzymatically active transpososome.

**Fig. 2.**
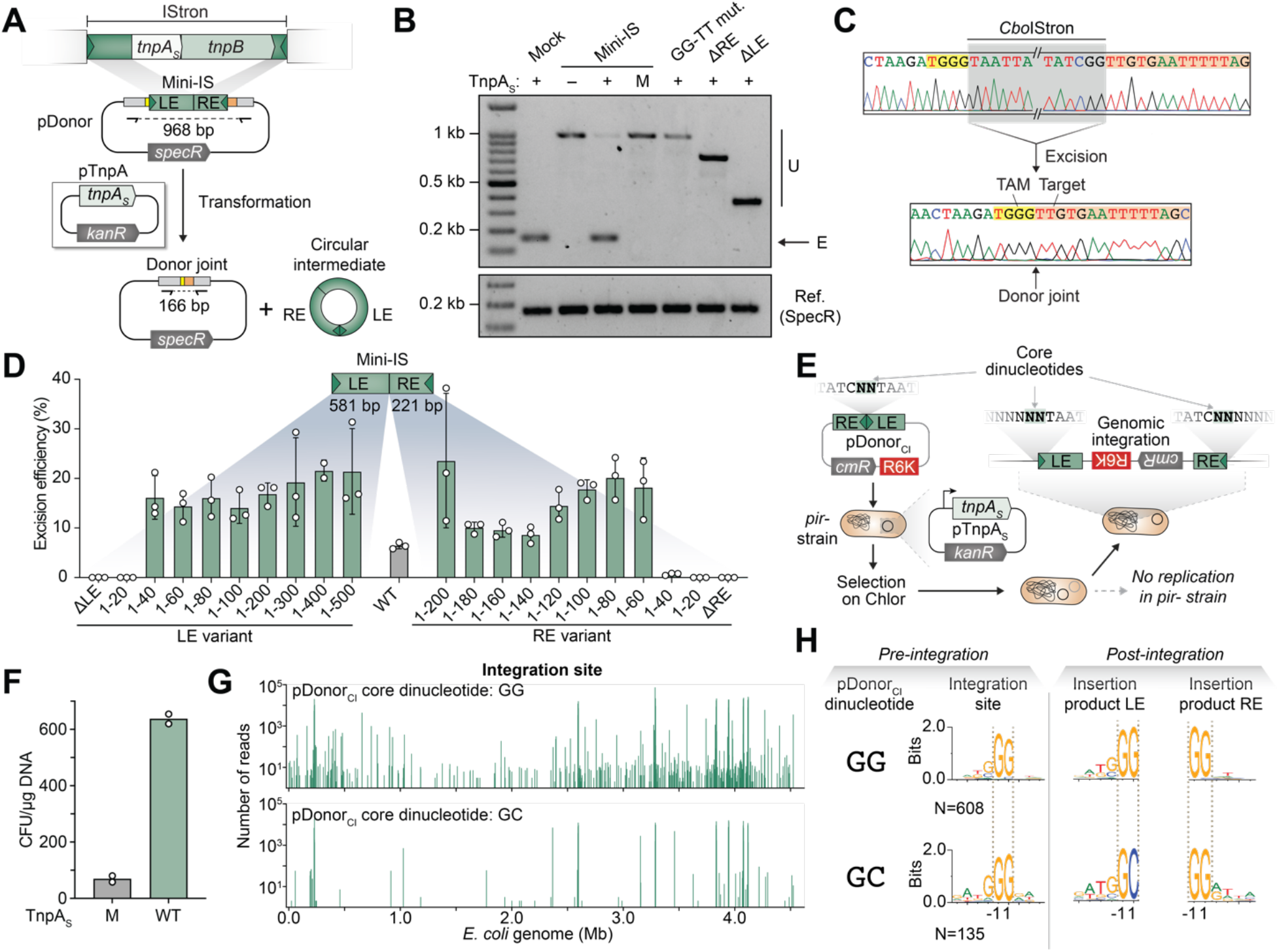
*Cbo*TnpA_S_ catalyzes efficient IStron excision and integration, with unique dinucleotide requirements. (**A**) Schematic of transposon excision assay using a *Cbo*TnpA_S_ expression plasmid (pTnpA_S_) and *Cbo*IStron donor plasmid harboring a minimal transposon (Mini-IS) with LE and RE boundaries (pDonor). Expected substrates and products generated upon transposon excision by PCR are indicated, as are the primer binding sites. (**B**) Gel electrophoresis and Sanger sequencing (**C**) of PCR products from **A**, demonstrating that TnpA_S_ is active in recognizing and excising the IStron. Cell lysates were tested after overnight expression of TnpA_S_ with the indicated substrates, which included an IStron mutant containing mismatched dinucleotides (LE: 5′-GG-3′, RE: 5′-TT-3′), and IStrons with RE or LE deletions. Mock denotes a positive excision control, U and E refer to unexcised and excised products. M denotes a S67A TnpA_S_ mutant. Sanger sequencing is shown at the right, with the rejoined TAM and putative ωRNA-matching target highlighted in yellow and orange, respectively. (**D**) Quantitative PCR-based assay to determine the minimal left end (LE) and right end (RE) sequences necessary for efficient IStron excision. Serial truncations were tested, starting with a WT substrate containing 581 bp and 221 bp derived from the native LE and RE, respectively. (**E**) Schematic of transposon integration assay using a TnpA_S_ expression plasmid (pTnpA_S_) and IStron circularized intermediate donor plasmid harboring abutted LE and RE sequences (pDonor_CI_). With this suicide vector that cannot propagate in a *pir–* strain, transposon integration events can be enriched using chloramphenicol selection and deep-sequenced using TagTn-seq (**Materials and Methods**). (**F**) Cell viability data from experiments in **E**, plotted as colony forming units (CFU), when cells contained either mutant S67A (M) or WT TnpA_S_. (**G**) Genome-wide distribution of TagTn-seq reads from experiments in **E** using WT TnpA_S_, mapped to the *E. coli* genome. Data are shown for pDonor_CI_ substrates containing either a GG (top) or GC (bottom) dinucleotide. (**H**) Meta-analyses of target site preferences and integration product dinucleotides at the LE and RE junctions, for the genome-wide insertion data with GG and GC dinucleotide substrates shown in **G**; the number of unique integration sites is indicated. The preferred genomic target motif is GG for both substrates, but high-throughput sequencing across the LE and RE junctions for integration products clearly reveals that non-canonical dinucleotides in pDonor_CI_ template correspond to non-canonical dinucleotides at the LE junction upon recombinational integration.

Having confirmed that TnpA_S_ transposition proceeds via a circular dsDNA intermediate, we decided to take a closer look into the integration reaction using a modified transposition assay. We expressed TnpA_S_ in the presence of non-replicative suicide vectors containing abutted transposon ends, thus mimicking the circularized intermediate (pDonor_CI_), such that genomic integration would lead to a selectable antibiotic resistance phenotype (**Fig. 2E**). After first demonstrating that cellular survival was dependent on catalytically active TnpA_S_ (**Fig. 2F**), we applied a next-generation sequencing (NGS) approach to interrogate genome-wide integration specificity (**fig. S6E**). These experiments revealed hundreds of unique genomic integration sites that were broadly distributed and shared a strict GG dinucleotide requirement (**Fig. 2G,H and fig. S6F,G**), with additional preferences upstream to yield a 5′-TGGG-3′ target site motif, in excellent agreement with the bioinformatically predicted TAM (**fig. S5C**). Target site selectivity for large serine recombinases can be modified by mutating core dinucleotides within the recombination sequence (*18*, *59*, *60*), and we wondered whether small TnpA_S_ recombinases would similarly switch their target site selectivity if the GG dinucleotide were altered. Intriguingly, TnpA_S_ still targeted 5′-TGGG-3′ sites when pDonor_CI_ contained a GC dinucleotide, but when we carefully inspected the actual genomic integration products from NGS data, we found that the LE and RE harbored non-matching GC and GG dinucleotides, respectively (**Fig. 2G,H**). Further testing of additional modified dinucleotides on pDonor_CI_ revealed an unexpected diversity of preferred integration sites and LE/RE genomic integration products (**fig. S6G**), highlighting the need for additional biochemical experiments to resolve the detailed mechanism of TnpA_S_-mediated recombination.

Beyond providing new insights into the transposition pathway, these experiments demonstrate that IS607 transposon loss is a direct consequence of TnpA_S_ activity, just as IS605 transposon loss is a consequence of TnpA_Y_ activity (*7*, *21*). Given the pervasive presence of TnpB nucleases in both transposon families, we hypothesized that RNA-guided DNA cleavage would be essential to avoid IS607 extinction, similar to its role in IS605 elements (*7*), thus representing a potentially universal function across diverse transposon families. We next set out to investigate *Cbo*TnpB and its impacts on *Cbo*IStron maintenance and spread.

### IStrons encode active TnpB nucleases that promote transposon maintenance and spread

We sought to first confirm a direct interaction between *Cbo*TnpB and its ωRNA, by performing RNA immunoprecipitation sequencing (RIP-seq; **Fig. 3A**). TnpB strongly enriched its expected substrate, but we were surprised to detect a strong shoulder in the read coverage located 42 nucleotides (nt) downstream of the start site for the ωRNA covariance model, suggesting the possibility of intramolecular processing (**Fig. 3B**). When we repeated the experiment using a TnpB variant containing an inactivating mutation in the RuvC nuclease domain (dTnpB), the shoulder was entirely abolished, implicating TnpB in enzymatic processing of the precursor transcript comprising its own mRNA fused directly to the ωRNA (**Fig. 3B**). Similar results were recently described for other TnpB homologs, suggesting a generalizable role of transposon-encoded nucleases in ωRNA biogenesis (*61*). Whether a similar feature is also true for related bacterial IscB nucleases and eukaryotic Fanzor nucleases remains to be determined.

**Fig. 3.**
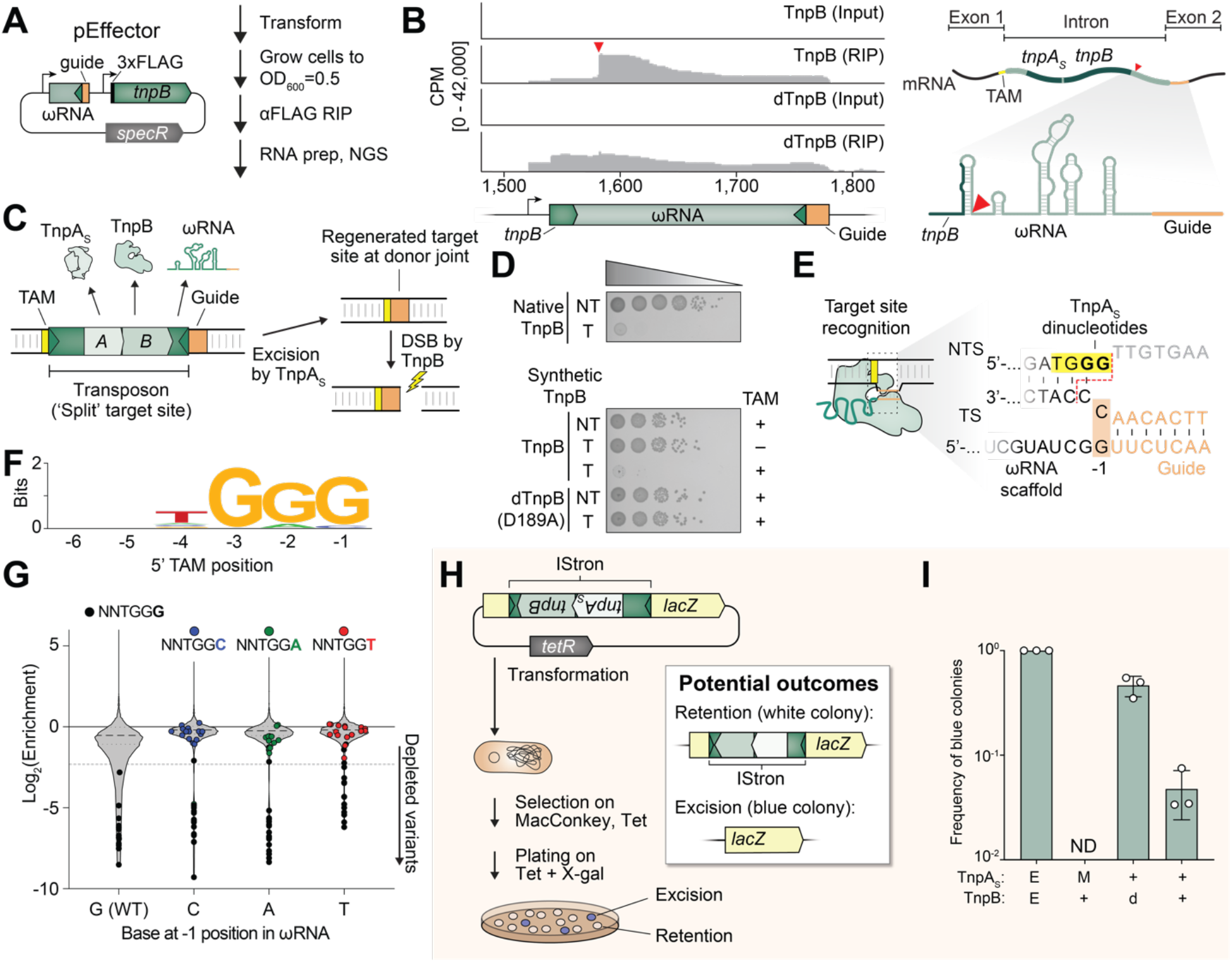
*Cbo*TnpB is a potent RNA-guided nuclease that prevents *Cbo*TnpA_S_-mediated transposon extinction. (**A**) Schematic of RIP-seq workflow to uncover RNA binding partners of *Cbo*TnpB using the pEffector shown. (**B**) RIP-seq read coverage for experiments with WT TnpB and RuvC-inactivated dTnpB (D189A) mapped to pEffector (left), alongside associated input controls. The pre-ωRNA processing site is indicated with a red triangle, in both the graph and the RNA schematic shown at the right. The green region labeled “*tnpB*” corresponds to the 3′ end of the ORF. (**C**) Schematic showing the regenerated target site that is produced upon transposon excision, with abutted TAM and target site. (**D**) Bacterial spot assays demonstrate that TnpB is highly active for RNA-guided DNA cleavage of the donor joint, as assessed by plasmid interference assays. Cells expressing TnpB from a native IStron or synthetic expression plasmid context were transformed with a target bearing (T) or no-target bearing (NT) plasmid, transformants were serially diluted, plated on selective media, and cultured at 37 °C for 24 hours. Additional controls included a mutant TAM (“–”, 5′-ACCC-3′) or RuvC-inactive (D189A) dTnpB. (E) Schematic indicating the uncertainty over whether nucleotides within the ωRNA scaffold might influence TAM specificity through direct base-pairing, especially since TnpA_S_ could theoretically recognize either of the two adjacent GG core dinucleotides defining the transposon boundary. NTS, non-target strand, TS, target strand. (**F**) Results from a TAM library cleavage assay using a wild-type ωRNA, revealing that *Cbo*TnpB requires a consensus 5′-TGGG-3′ TAM for efficient DNA cleavage. The WebLogo was generated using the 20-most depleted sequences after deep sequencing pTarget from surviving colonies (see **fig. S7**). (**G**) Violin plots showing enrichment of TAM sequences from TAM library assays using variant TnpB-ωRNA expression plasmids with the indicated nucleotide in the –1 position of the ωRNA. Data are plotted as the log_2_-fold enrichment relative to the input library, with specific members highlighted; dotted line represents 5-fold depletion. All ωRNA variants depleted only 5′-TGGG-3′ TAMs, indicating an absence of base-pairing at the –1 position and supporting the idea that this is the last nucleotide of the ωRNA scaffold. (**H**) Schematic of the assay to measure transposon fate in *E. coli* with TnpA_S_ and TnpB– ωRNA, and (**I**) bar graph showing the frequency of transposon excision/retention for each condition, quantified by blue/white colony screening. *Cbo*IStron variants (mutant S67A TnpA (M), dTnpB (D189A) or WT) were inserted at a compatible TAM in a plasmid-encoded *lacZ* and used for transformation of a *lacZ–* strain. White or blue colonies indicate transposon retention or excision, respectively. Data are mean ± s.d. (n = 3); E, empty vector; ND, not detected.

Based on the prior work with IS605-family TnpB homologs (*7*, *8*), we hypothesized that *Cbo*TnpB would recognize and cleave the scarless donor joint generated upon TnpA_S_-mediated IStron excision, in which the TAM and guide-matching target region are directly adjoined (**Fig. 3C**). Using a plasmid interference assay, in which successful targeting results in cell lethality under antibiotic selection, we found that TnpB was highly active for RNA-guided DNA cleavage, causing a ∼10^5^-fold decrease in colony-forming units (**Fig. 3D**). This reaction strictly required guide RNA–target DNA complementarity, a cognate TAM, and an intact RuvC active site, and was similarly effective whether we used a synthetic expression vector with separate promoters for *tnpB* and ωRNA or a native-like *Cbo*IStron (lacking *tnpA*) with single transcript-derived *tnpB*-ωRNA (**Fig. 3D**).

Our transposon insertion assays revealed that TnpA_S_ selects 5′-TGGG-3′ target sites, with only the GG dinucleotide being strictly required. We were initially uncertain where to define the transposon boundary for integrated elements, which affected our ability to confidently determine the precise TAM required by TnpB during DNA recognition and cleavage (**Fig. 3E**). We thus performed an unbiased cellular TAM library assay using a target plasmid containing a degenerate 6N library, in which cleaved molecules are strongly depleted under selective growth conditions (**fig. S7A**). These experiments revealed a robust 5′-TGGG-3′ TAM motif (**Fig. 3F**) that precisely matched the target site recognized by TnpA_S_ during transposition (**Fig. 2H**), demonstrating that, as with TnpA_Y_/TnpB in IS605 elements (*7*), distinct transposase and nuclease enzymes have converged to specify the same DNA motif. Interestingly, the guanine at the −3 position was most strictly discriminated, explaining how the related 5′-T**<u>C</u>**GG-3′ motif adjacent to the target site within the transposon itself would be avoided for self-cleavage (**fig. S7B**). We observed that the last two nucleotides in the ωRNA scaffold sequence matches the last nucleotides of the TAM, raising a question whether our assignment of the guide and scaffold was correct. We decided to test if the nucleotide at the −1 position in ωRNA was recognizing the target through base-pairing (**Fig. 3E**) by repeating TAM library assays using modified ωRNAs, in which the −1 position was replaced with all other possible nucleotides. We found that all modified ωRNAs were active for cleavage and in all cases, the only TAMs that were consistently depleted bore the 5′-TGGG-3′ sequence, irrespective of the nucleotide present at −1 position, showing no evidence of base-pairing (**Fig. 3G and fig. S7C–E**). Altogether, these results confirm that *Cbo*TnpB recognizes 5′-TGGG-3′ TAM, and that the ωRNA scaffold ends with 5′-…UCGG-3′, in agreement with a recent report for another IS607-family TnpB (*12*).

We recently demonstrated that TnpB is essential to preventing permanent loss and extinction of IS605-family transposons, an inherent risk due to the excision and scarless rejoining activity of the TnpA_Y_ transposase. By using ωRNA guides to target and cleave the empty donor joints generated upon IS605 transposon excision, TnpB nucleases trigger efficient recombination with homologous chromosomes, thereby reinstalling transposon copies and promoting homing into new target sites, through a mechanism we termed peel-and-paste / cut-and-copy (*7*). IS607-family transposons are similarly at risk of permanent loss as a consequence of the transposition mechanism catalyzed by TnpA_S_, despite mobilizing through a dsDNA intermediate (**Fig. 1A**). Given these commonalities, as well as the convergent recognition of the same target motif and TAM by *Cbo*TnpA_S_ and *Cbo*TnpB, we hypothesized that TnpB nucleases are also essential to prevent transposon extinction of IS607-family elements.

We set out to test this hypothesis for the *Cbo*IStron model system, by applying our recently developed assays to monitor both transposon loss and DSB-mediated transposon homing (*7*). First, we inserted the native *Cbo*IStron into *lacZ* (white colonies) in the anti-sense orientation, to avoid confounding effects of splicing, and monitored transposon excision via regeneration of a *lacZ*+ phenotype (blue colonies), in the presence of WT/mutant TnpA_S_ and TnpB (**Fig. 3H**). TnpA_S_ drove extensive and rapid excisional transposon loss, leading to 47% of colonies regaining a blue phenotype, and this frequency was suppressed 10-fold in the presence of WT but not nuclease-dead TnpB (**Fig. 3I**). The role of TnpB in preserving transposon copies was further corroborated by diagnostic PCR-based genotyping (**fig. S8A**). To prove that transposon maintenance was the consequence of donor joint cleavage followed by homologous recombination, we next monitored the ability of TnpB-ωRNA to specifically mobilize the *Cbo*IStron into empty donor joints present in the *E. coli lacZ* locus (**fig. S8B**). The presence of WT TnpB nuclease strongly reduced the overall transformation efficiency, as expected, and a significant proportion (>80%) of surviving colonies indeed represented faithful recombination products, as assayed by exhibiting a white-colony phenotype (**fig. S8C**). This effect was completely ablated by nuclease-dead TnpB, confirming the critical role of TnpB-induced DSBs in the process.

Collectively, these experiments powerfully expand our paradigm for the essential role of TnpB in promoting transposon survival to one of the most pervasive transposon families found in nature, IS607-family elements.

### Group I intron splicing levels are regulated by TnpB and ωRNA structure

Compared to simpler IS605- and IS607-family transposons, IStrons are unique in their predicted ability to self-splice at the RNA level and thereby rejoin flanking exonic segments, which would presumably render their presence – at the DNA level – phenotypically silent. The *Cbo*IStron displays all of the expected secondary structural elements for group I introns (*46*, *47*), and our analyses tentatively predicted the 5′SS and 3′SS as perfectly aligning with the transposon LE and RE boundaries, such that the DNA donor joint would match the spliced mRNA exon-exon joint (**Fig. 1B**). However, we set out to empirically investigate splicing by developing a cellular assay in which unspliced and spliced products are directly detected by RT-PCR (**Fig. 4A**). Using a minimal IStron substrate that lacked both *tnpA* and *tnpB* ORFs, we readily detected the dual presence of both species, and moreover, the spliced product exhibited scarless joining of the two flanking exons (**Fig. 4B**), with the exact same nucleotide connectivity as seen with the DNA donor joint upon TnpA_S_-mediated excision (**Fig. 2C, 4C**). Splicing was abolished by removing the P7-P9 loop region that comprises the intron active site (**fig. S5D**) or by introducing point mutations at the 5′SS (**Fig. 4B**). We also observed the exact same splicing products from *in vitro* biochemical reactions, indicating that no protein factors were required (**fig. S9A**).

**Fig. 4.**
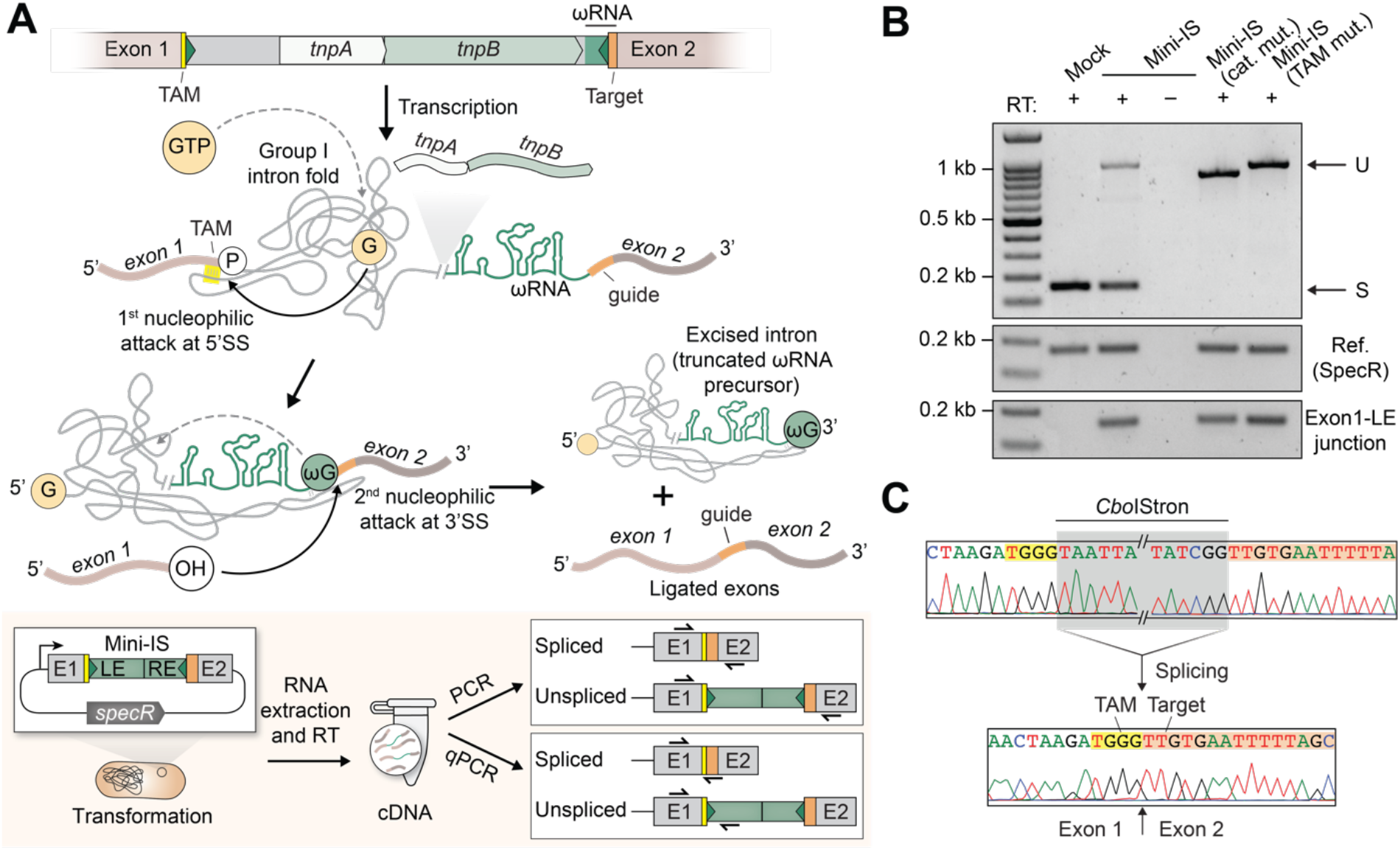
*Cbo*IStrons encode functional self-splicing ribozymes that regenerate transposon-free transcripts. (**A**) Schematic of general IStron splicing mechanism (top) and *E. coli*-based cellular splicing assay (bottom). Exogenous GTP binding by the folded group I intron leads to a transesterification reaction at the 5′ splice site (5′SS), followed by attack of the 3′SS by exon 1 to yield the ligated exon-exon product and excised intron. Spliced/unspliced products are detected and/or quantified by RT-PCR and RT-qPCR, respectively, using the primer pair strategies indicated at the bottom. (**B**) Agarose gel electrophoresis of RT-PCR products from splicing assays in **A** with the indicated constructs, which shows the extent of unspliced (U) and spliced (S) products (top) relative to reference amplicons for a SpecR drug marker (middle) and exon1–LE junction (bottom). RT, reverse-transcriptase; Mock denotes a positive splicing control; IStron (cat. mut.) contains a P7-P9 loop deletion in the intron catalytic core; IStron (TAM mut.) contains 5′-TGTA-3′ in the TAM and thereby disrupts base-pairing required for 5′ SS recognition. (**C**) Sanger sequencing of RT-PCR products from **B**, for both the unspliced exon-intron boundaries (top) and the spliced exon-exon product (bottom). These sequences are identical to the nucleotide sequences of unexcised and excised DNA sequences shown in Fig. 2C.

Unspliced transcripts produced from native *Cbo*IStrons presumably satisfy numerous functions, of which some are mutually compatible, but others are mutually exclusive. Both unspliced and spliced introns can serve as protein-coding mRNAs for TnpA_S_ and TnpB expression, but spliced introns can no longer act as functional ωRNAs since the splicing reaction severs the ωRNA scaffold from the ωRNA guide region (**Fig. 4A**). Additionally, TnpB-mediated ωRNA processing (**Fig. 3B**) would segregate the 5′SS and 3′SS onto two distinct molecules, rendering only trans-splicing possible, and TnpB-ωRNA binding would also likely obstruct physical access to the 3′SS. We therefore hypothesized that the structure of the intron RNA 3′ end (**Fig. 5A**), the presence of TnpB, or both, would regulate the relative efficiencies of splicing and RNA-guided DNA cleavage, and set out to test this hypothesis via targeted perturbations.

**Fig. 5.**
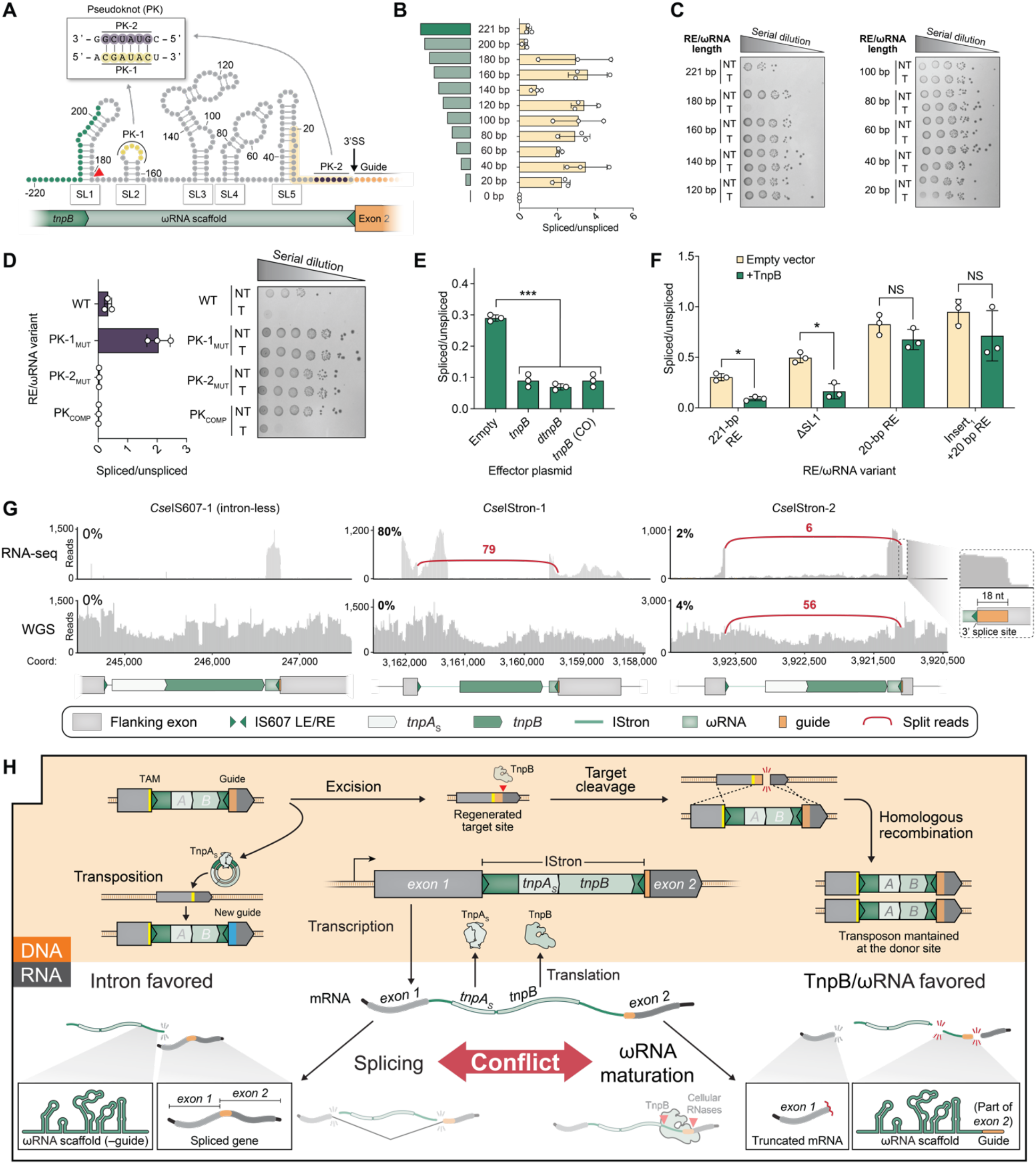
Competition between intron splicing and TnpB–ωRNA activity establishes a balance between transposon stealth and preservation. (**A**) Schematic of *Cbo*IStron ωRNA secondary structure encoded within the transposon RE, with stem-loops (SL), truncation coordinates, and pseudoknot (PK) motifs labeled. PK-2 overlaps with nucleotides important for 3′ splice site (3′SS) recognition. Red arrow, TnpB processing site; green dots, *tnpB* ORF; yellow shading, minimal sequence required for efficient splicing. (**B**) RT-qPCR analysis of splicing efficiency for IStron variants in which the RE/ωRNA region was systematically truncated relative to the full-length construct (221 bp). The large splicing change with the 180-bp construct suggests sequence and/or structural features around this position that repress splicing in the full-length design. (**C**) Bacterial spot assays for the same RE/ωRNA deletion constructs in **B**, in which RNA-guided DNA cleavage leads to cell death. Cells expressing TnpB were transformed with either a target bearing (T) or no target bearing (NT) plasmid, and transformants were serially diluted, plated on selective media, and cultured at 37 °C for 24 hours. A 200-bp construct was not tested due to the beginning of ωRNA covariation model overlapping with the *tnpB* ORF. (**D**) RT-qPCR analysis of splicing efficiency (left), and spot assays to monitor RNA-guided DNA cleavage activity (right), for the indicated RE/ωRNA pseudo-knot mutations, plotted as in **B** and **C**. PK-1_MUT_ and PK-2_MUT_ contain mutations to either motif, whereas PK_COMP_ contains compensatory mutations in both motifs. The results indicate that ωRNA PK disruption abrogates TnpB-mediated DNA cleavage, while any mutation to the downstream PK motif (PK-2) abrogates intron splicing; intron splicing is strongly stimulated by mutations to the upstream PK motif (PK-1). (**E**) RT-qPCR analysis of splicing efficiency in the presence of a second effector plasmid harboring *tnpB*, *dtnpB*, or a codon-optimized (CO) *tnpB* gene, revealing the repressive role of TnpB. Empty refers to an empty vector control. Statistical significance was determined using Welch’s t-test, *** denotes p-value <0.001. (F) RT-qPCR analysis of splicing efficiency in the absence/presence of TnpB for the indicated RE/ωRNA variants. The repressive effect of TnpB on splicing is largely ablated when the ωRNA scaffold is missing (20-bp RE) or replaced with an unrelated sequence (Insert_1_+20-bp RE). Statistical significance was determined using Welch’s t-test, * denotes p-value <0.05, NS, not significant. (**G**) RNA-seq and whole-genome sequencing (WGS) data from three representative IS607 elements in *C. senegalense*, one without an intron (left) and two that encode identifiable group I introns (middle, right). The number and connectivity of spliced exon-exon junction reads are indicated (red), and quantitative comparison of exon-intron and exon-exon reads yields an apparent splicing percentage at the RNA level (RNA-seq graphs, top left); similar measurements were also made at the DNA level (WGS graphs, top left). These analyses indicate the *Cse*IStron-1 undergoes highly efficient splicing without any evidence of transposon excision, whereas the inefficient *Cse*IStron-2 splicing (2%) can be fully explained by low-level transposon excision within the bacterial culture, as reported by WGS. High versus low splicing correlates with low versus high ωRNA coverage for *Cse*IStron-1 and CseIStron-2, respectively. The inset shows overlap between the ωRNA and 3′SS. (**H**) Overall model for the balanced effects of intron splicing, TnpB– ωRNA, and TnpA_S_ transposition activity in the maintenance and spread of IS607-family IStron elements. Similarly to IS200/IS605-family transposons, scarless DNA excision by TnpA_S_ for IS607-family elements leads to transposon loss at the donor site and thus eventual transposon extinction, without the crucial function provided by TnpB–ωRNA in generating targeted DNA double-strand breaks and triggering homologous recombination to maintain presence of the transposon (top). Unlike canonical IS605- and IS607-family transposons, group I intron-containing IStrons mitigate their fitness costs on the host by splicing themselves out of interrupted transcripts at the RNA level, thereby restoring functional gene expression (bottom, middle). Splicing and ωRNA maturation are mutually exclusive, since splicing severs the ωRNA scaffold and guide sequences, and TnpB represses splicing through competitive binding of the 3′SS. The competition between intron splicing and TnpB–ωRNA activity thus serves to regulate the dual objectives of maintaining transposon stealth and promoting transposon proliferation for IStron elements. A similar mechanism is hypothesized for IS200/IS605-family IStrons.

Any deletions to the intron structural core overlapping the *Cbo*IStron LE were deleterious for splicing, as expected (**fig. S9B**). When we introduced serial deletions within the RE, starting from the ωRNA 5′ end, we found that splicing was dramatically increased for most constructs that eroded key stem-loops (SL), most of which abolished RNA-guided DNA cleavage activity by TnpB (**Fig. 5B,C and fig. S9B,C**). Targeted SL deletions, alone or in combinations, again demonstrated that most ωRNA disruptions were actually stimulatory for splicing while ablating TnpB activity, with the exception of SL5, which was essential for splicing (**fig. S9D,E**); further deletion experiments confirmed that sequences within SL5 are essential and likely directly participate in the splicing reaction (**fig. S9F**). Lastly, we inserted three arbitrary *lacZ*-derived sequences upstream of a minimal 20-bp RE/ωRNA to complement the native ωRNA length and found that these constructs were more active for splicing than the WT *Cbo*IStron, similarly to the minimal splicing variant (**fig. S9G**). These results supported the notion that splicing and TnpB-ωRNA activity are inversely correlated and that RE/ωRNA truncations stimulate splicing in a TnpB-independent manner, indicating that RNA structure alone influence splicing activity.

We were particularly intrigued by a predicted pseudoknot (PK) interaction within the ωRNA that is widespread across diverse TnpB and Cas12 homologs (*54–58*) (**fig. S10A**), and which would sterically occlude nucleotides required for 3′SS recognition (**Fig. 5A**). PK formation was essential for TnpB activity, as DNA cleavage was eliminated with mutations to either PK-1 or PK-2, but was restored with compensatory mutations to both motifs (**Fig. 5D**). When we investigated splicing, we were surprised to find that selective mutations to just the upstream PK-1 motif, which would disrupt PK formation without affecting the 3′SS, increased the splicing ratio 7-fold, presumably by preventing formation of splicing-incompetent RNA conformations (**Fig. 5D**). Conversely, mutations to the downstream PK-2 motif, or compensatory mutations to both PK-1 and PK-2, eliminated splicing because of the overlap between these mutations and the 3′SS.

To test whether TnpB also directly impacted splicing, we designed a modified assay in which full-length TnpB was expressed *in trans* relative to the mini-*Cbo*IStron construct, to avoid size-dependent splicing effects (**fig. S10B-D**). We found that TnpB potently reduced splicing efficiency regardless of whether it possessed an intact nuclease active site or not (**Fig. 5E**), indicating that ωRNA binding alone, with or without processing, had a repressive effect. To confirm that this observation was dependent on specific ωRNA structural features, we repeated these experiments with constructs that harbored various sequence perturbations to the RE/ωRNA sequence. Splicing was only repressed in a TnpB-dependent manner when the majority of the ωRNA scaffold was present within the intron, and this repression was lost when the full-length ωRNA was replaced with a *lacZ*-derived sequence insertion (**Fig. 5F**). Collectively, these data indicate that TnpB, as well as RNA-intrinsic properties, regulate the efficiency of intron splicing.

Finally, we sought native *in vivo* data supporting the antagonistic relationship between IStron-encoded introns and ωRNAs. Published sequencing data were unavailable for *Clostridium botulinum* strains that encode IS607-family *Cbo*IStrons, so we turned our attention to an available *Clostridium senegalense* strain (ATCC 25772) that harbors two IS607-family IStrons and 9 additional intron-lacking IS607 elements (**fig. S11A, table S1**). After culturing the strain, isolating total RNA, and performing RNA-sequencing (RNA-seq), we observed dramatic differences in splicing between the two *Cse*IStron loci, with IStron-1 exhibiting 80% splicing efficiency versus only ∼2% for IStron-2 (**Fig. 5G**); in both cases, the 5′SS and 3′SS precisely corroborated our predictions based on comparative genomics and transposon LE/RE boundary assignments. Remarkably, we observed prominent ωRNA coverage for *Cse*IStron-2 and intron-less IS607 elements, with the expected ∼18-nt guide sequence extending beyond the RE boundary, whereas ωRNA reads were largely absent from the *Cse*IStron-1 locus for which splicing was highly efficient (**Fig. 5G and fig. S11B**). There was also an abundance of RNA-seq read pairs that terminated at the *Cse*IStron-2 5′SS, indicative of prematurely aborted splicing reaction products that only progressed through the initial cleavage at the upstream exon-intron junction (**fig. S11C**).

IStrons yield the exact same nucleotide connectivity after RNA splicing and DNA excision (**Fig. 2C,4C**). To determine whether splicing calls from RNA-seq data were simply the result of low-level, heterogeneous transposon DNA excision, we performed whole-genome sequencing (WGS) for the same *C. senegalense* cultures. Strikingly, we observed a similar frequency of *Cse*IStron-2 excision (and thus apparent ‘splicing’) from the WGS data (**Fig. 5G**), a conclusion that was corroborated by additional validation from targeted deep sequencing (**fig. S11D**). In contrast, no evidence of DNA-level loss for *Cse*IStron-1 or an intron-less IS607 element was observed. These data further strengthen our conclusion that group I intron splicing and ωRNA biogenesis are negatively correlated, while also revealing the mutable nature of IS transposons on short timescales.

## DISCUSSION

This work reveals an intricate balancing act that IStrons evolved to satisfy mutually exclusive functions of transposon-encoded non-coding RNAs (**Fig. 5H**). Efficient self-splicing of the group I intron is crucial for these elements to access a larger targetable space in the genome, since transposon insertions within coding genes, even those essential to the host, can occur without significant phenotypic consequences upon exon-exon ligation; this conclusion is readily borne out by the genetic context of diverse IStrons found in Clostridial genomes (*31–35*) (**Fig. 1C and fig. S3C,S5B**). Yet the intron 3′ end also contains the ωRNA scaffold whereas the downstream exon contains the critical targeting sequence, such that splicing effectively severs the head (guide) from the body (scaffold) of the guide RNA (**Fig. 5H**). And without a functional ωRNA, TnpB is unable to maintain its function in transposon maintenance through targeted cleavage of empty donor joints upon TnpA-mediated transposon excision. Thus, IStron transcripts must exhibit the ability to exist in both splicing-competent and ωRNA-competent states, with the relative proportion dictating the balance between fitness cost maintenance (RNA-driven intron splicing) and selfish transposon preservation (RNA-guided DNA cleavage).

Our results highlight the separate roles of TnpB and RNA structure in establishing this sensitive equilibrium. TnpB-ωRNA binding acts to physically obstruct the second step of splicing (**Fig. 5E,F**), preventing the upstream exon from accessing the scissile phosphate at the 3′ intron-exon boundary through steric occlusion. The RNA alone also regulates its own fate through a critical pseudoknot interaction within the ωRNA, which competes for direct binding to the 3′ splice site and thereby represses splicing (**Fig. 5A,D**). By systematically mutagenizing the RE/ωRNA region of the *Cbo*IStron and studying the effects on splicing and TnpB activity, we were able to provide evidence for these conflicting conformational states, and *in vivo* data from native IStron elements in *C. senegalense* further corroborate the inverse relationship between intron splicing and ωRNA biogenesis (**Fig. 5G**). However, future structural, mutational, and chemical probing studies will be needed to directly interrogate structural dynamics of the non-coding RNA, and to investigate their ramifications on long-term transposon proliferation in both native and engineered settings.

On the surface, co-occurrence of group I introns with IS605- and IS607-family transposons appears ill-suited to satisfy their selfish needs, since autonomous functions of the intron RNA are subverted via the conflict with ωRNA. The well-studied co-option of homing endonucleases (HEs) also provides a seemingly effective dispersion strategy for group I introns (*28–30*), since these DNA endonucleases can be encoded and expressed from within the intron without disrupting access to the 5′ or 3′ splice sites (5′/3′SS). But in fact, IStrons provide far greater flexibility in mobilization, since the association with diverse tyrosine-family (TnpA_Y_) and serine-family (TnpA_S_) recombinases ensures even greater opportunities for broad intron dissemination (**Fig. 1A,2**), without being limited to homologous sites targeted by HEs. Moreover, the programmable nature of TnpB nucleases not only enables homing to each new insertion site, but also addicts the cell to maintaining the intron at its donor site (**Fig. 3, 5H**). This capability is uniquely enabled by the elegant overlap between the transposon LE/RE boundaries at the DNA level, and the intron 5′SS/3′SS boundaries at the RNA level, ensuring a perfect correspondence between the nucleotide sequence inserted during DNA transposition and removed again during RNA splicing (**Fig. 2C,4C**). Our comprehensive bioinformatics survey of group I introns provides a compelling view of the diverse strategies that these ancient ribozymes have adopted over evolutionary timescales (**Fig. 1D**), and may reveal novel genetic associations with additional enzymatic modules in the future. Interestingly, transposons are also capable of mediating the spread of spliceosomal introns in eukaryotes (*62*, *63*), suggesting that the synergy of transposition and splicing spans the cross-kingdom boundary.

Another key finding of this study relates to the central role of TnpB in IS607 transposon preservation. Earlier work from the Zhang and Siksnys labs identified TnpB as an RNA-guided DNA endonuclease (*8*, *9*), and our recent study was the first to prove that this biochemical activity promotes selfish spread of IS605-family transposons by triggering a composite transposition pathway, together with the TnpA_Y_ transposase, that we termed peel-and-paste, cut-and-copy (*7*). Here, we demonstrate that TnpB plays an equally crucial function in promoting the spread of IS607-family transposons (**Fig. 3H,I and fig. S8**), one of nature’s most ubiquitous selfish genetic element found in all three domains of life (*10*, *15*, *64*). IS605 and IS607 transposons encode phylogenetically distinct transposases and mobilize through different intermediates, but they both suffer the same risk of permanent loss due to their scarless mechanism of excision that is uncoupled from integration (**Fig. 1A**). Remarkably, both transposon families encode structurally similar ωRNA guides that overlap the transposon right end (RE), despite requiring vastly different DNA sequences/structures to satisfy transposase-transposon recognition constraints (**Fig. 1C and fig. S5E,F**). This flexibility, together with the convergent specificity for overlapping target and transposon-adjacent motifs during DNA integration and DNA cleavage by TnpB and TnpA, respectively (**Fig. 2G-H, 3C-G)**, has ensured functionally intertwined coupling of enzymatic machineries that catalyze transposon spread and transposon retention. It is noteworthy that TnpB evolved disparate preferences for TA-rich and G-rich TAMs during its evolution with IS605 and IS607 transposon, respectively, likely predating the later PAM diversification that would arise for Cas12 nucleases during the evolution of antiviral adaptive immunity (*65–69*).

Insertion sequence (IS) transposons were initially defined as short DNA segments encoding only the enzymes necessary for their transposition (*3*). IStrons provide a striking counterexample, showcasing the multiple overlapping biochemical reactions that all contribute to the complex behavior of a transposable element, with as much attention focused on DNA preservation and phenotypic silencing as on transposition itself. In turn, IStrons and IS605/IS607-family transposons, more generally, also highlight the unique molecular functionalities that have arisen during co-evolution of distinct machineries encoded within mobile genetic elements. It is via IS607-family transposons that TnpB nucleases likely gained access to the eukaryotic domain, giving rise to homologous proteins called Fanzors, with viruses likely serving as a potent mode of transmission (*10–13*). And in the ensuing diversification of eukaryotic Fanzors, co-evolutionary relationships have yielded fascinating associations with new molecular actors, including *piggyBac*, *Tc1/mariner*, and *Helitron* transposases, alongside other, as yet uncharacterized genes (*10*, *13*). It appears likely that new biochemical activities and biological functions will continue to be uncovered within this fascinating context.

## MATERIALS AND METHODS

### Protein detection and database curation

#### TnpB

The TnpB database was previously curated (*7*). As described therein, homologs were comprehensively detected using the *H. pylori* TnpB (*Hpy*TnpB) amino acid (aa) sequence (NCBI Accession: WP_078217163.1) and the *G. stearothermophilus* TnpB amino acid sequence (NCBI Accession: WP_047817673.1) as seed queries for two independent iterative JackHMMER (HMMER suite v3.3.2) searches against the NR database (retrieved on 06/11/2021), with an inclusion and reporting threshold of 1e-30. The union of the two searches were taken, and proteins that were less than 250 aa were removed to trim partial or fragmented sequences, resulting in a database of 95,731 non-redundant TnpB homologs. Contigs of all putative *tnpB* loci were retrieved from NCBI for downstream analysis using the Bio.Entrez package.

#### TnpA_Y_ and TnpA_S_

For *tnpB*-associated contigs, *tnpA*_Y_ was detected using the Pfam Y1_Tnp (PF01797) model for a HMMsearch from the HMMR suite (v3.3.2), with an E-value threshold of 1e-4. This search was performed on the curated CDSs of each contig from NCBI. IS elements that encoded *tnpB* within 1-kb of a detected *tnpA*_Y_ were defined as autonomous. Analysis of *tnpA*_S_ association with *tnpB* was performed with the same methodology mentioned above, but using the serine resolvase Pfam model (PF00239).

#### Arc-like ORF

A manually identified Arc-like protein (NCBI Accession: WP_003367503.1) was used as the seed query in a two-round PSI-BLAST search against the NR database (retrieved on 08/17/23). A neighborhood analysis was conducted on ORFs within 10 kb of all detected Arc-like ORF loci using HMMscan from the HMMR suite (v3.3.2) with the Pfam database of HMMs (retrieved on on 06/29/2021), and TnpB homologs were specifically identified using the TnpB-specific models produced from the JackHMMER performed in *Meers et al.* (*7*). High frequency associations with Arc-like ORFs were manually inspected, and putative functional associations were manually annotated.

### ncRNA covariation analyses

#### Group I Intron

The initial search for group I introns associated with *tnpB* was performed using models of available subclasses from the Group I Intron Sequence and Structure Database (GISSD), refined by Zhou *et al.* (*46*) and Nawrocki *et al.* (*70*). The 14 group I intron subclass models were searched against all identified *tnpB*-associated contigs with cmscan (Infernal v1.1.4). A liberal minimum bit score of 15 was used in an attempt to capture distant or degraded introns, and the identification of a putative IStron was supported by its proximity, orientation, and relative location to the nearest identified *tnpB* ORF. The remaining intron hits were considered associated with *tnpB* if they were upstream, on the same strand, and within 1 kb of a *tnpB* ORF. After manually inspecting the database of models and the boundaries of hits, only the catalytic subdomains of the intron were captured, resulting in the poor identification of other substructures both upstream and downstream of the hit. To address this, the boundaries of the group I intron found to be associated with *tnpB* genes were refined and used to generate a more accurate, comprehensive covariance model. A new model was built using a set of 103 *tnpB* loci, including the *C. botulinum* IStron experimentally tested in this study, and closely related loci. To build the new model, 1.5 kb upstream of *tnpB* gene were extracted and clustered by 99% length coverage and 99% alignment coverage using CD-HIT (*71*) (v4.8.1), to remove identical sequences. The resulting sequences were aligned using MAFFT (*72*) (v7.508) with the E-INS-I method for 8 iterations. The 5′ boundary of the intron (and LE of the IStron) was manually identified as the position of significant drop-off of sequence identity in the alignment; sequences were subsequently trimmed to that boundary. A structure-based multiple alignment was then performed using mLocARNA (*73*) (v1.9.1), with the following parameters:

*--max-diff-am 25 --max-diff 60 --min-prob 0.01 --indel −50 --indel-open −750 --plfold-span 100 --alifold-consensus-dp*

The resulting alignment with structural information was used to generate a new group I intron covariance model with the Infernal suite and verified/refined with R-scape at an E-value threshold of 1e-5. The resulting covariance model was used with cmsearch to discover new group I introns within our curated *tnpB*-associated contig database. The resulting sequences were aligned to generate a new CM model that was used to again search our *tnpB*-associated contig database. After refinement, the final group I intron CM model was searched against the entire NT database (retrieved on 08/29/23) with a higher bit-score of 40.

##### ωRNA

The initial boundaries of the ωRNA associated with IStron *tnpB* genes were identified in Meers *et al.* (*7*). To refine these models so that structures more representative of both IS605- and IS607-family IStrons, we extracted sequences 200-bp downstream and 50-bp upstream of the last nucleotide of *tnpB* ORFs, to define the RE and transposon boundaries. The ∼250-bp sequences were clustered by 99% length coverage and 99% alignment coverage using CD-HIT, to remove duplicates. The remaining sequences were then clustered again by 95% length coverage and 95% alignment coverage using CD-HIT. This was done to identify clusters of sequences that were closely related but not identical, as expected of IS elements that have recently mobilized to new locations. For the 100 largest clusters, which all had a minimum of 10 sequences, MUSCLE (*74*) (v3.8.1551) was used to align each cluster of sequences with default parameters. Then, each cluster alignment was manually inspected for the boundary between high and low conservation, or where there was a stark drop-off in mean pairwise identity over all sequences. This coordinate was annotated for each cluster as the putative 3′ end of the IS element. If there was no conservation boundary, sequences in these clusters were expanded by another 150 bp, in order to capture the transposon boundaries, and realigned. The consensus sequence of each alignment (defined by a 50% identity threshold up until the putative 3′ end) was extracted, and rare insertions that introduced gaps in the consensus were manually removed. With the 3′ boundary of the IS element, and thus the 3′ boundary of the *tnpB* ωRNA properly defined, a covariance model of the ωRNA for any *tnpB* clade of interest could now be built.

A 200-bp window of sequence upstream of the 3′ end for elements of the *Cbo*IStron clade and the *Cdi*IStron clade was extracted. A structurally-based multiple alignment was then performed using CMfinder (*75*) (v0.4.1.9) with the following parameters:

*-skipClustalw -minCandScoreInFinal 10 -combine -fragmentary -commaSepEmFlags x--filter-non-frag,--max-degen-per-hit,2,--max-degen-flanking-nucs,7,--degen-keep,--amaa*

The resulting structural motifs were then used to generate either a *Cbo*IStron- or *Cdi*IStron-specific ωRNA covariance model with Infernal and refined/verified with R-scape (*76*) at an E-value threshold of 1e-5. Each model was used to search against our *tnpB*-associated contig database with cmsearch at a bit-score of 40 to identify all structurally related ωRNA. ωRNA hits were considered associated with *tnpB* if they were downstream, on the same strand, and within 200 bp of a *tnpB* ORF. The alignment of hits from this search were refined/verified and visualized with R-scape.

### Phylogenetic analyses

#### TnpB and Arc-like ORF

For *tnpB* genes found in putative IStron elements, protein sequences were clustered at 95% length coverage and 95% alignment coverage using CD-HIT. The clustered representatives were taken and aligned using MAFFT with the E-INS-I method for 16 rounds. Post-alignment cleaning consisted of using trimAl (*77*) (v1.4.rev15) to remove columns containing more than 99% of gaps and manual inspection. The phylogenetic tree was created using IQ-Tree 2 (*78*) (v2.2.3) with a model of substitution identified using ModelFinder (*79*) (JTTDCMut+F+R10), and optimized trees with nearest neighbor interchange to minimize model violations. Branch support was evaluated with 1000 replicates of SH-aLRT, aBayes, and ultrafast bootstrap support (*80*) from the IQ-Tree 2 package. The tree with the highest maximum-likelihood was used as the reconstruction of the IStron *tnpB* phylogeny.

All Arc-like ORF hits were aligned using MAFFT with the E-INS-I method for 8 rounds. The rest of the analysis was identically performed as above. ModelFinder identified LG+I+R5 as the best-fit model.

#### Group I Introns

After the search of the NT database using an updated covariation model, hits smaller than 300 bp were removed. The remaining sequences were clustered at 90% length coverage and 90% alignment coverage using CD-HIT. The clustered representatives were taken and aligned using MAFFT with the E-INS-I method for 2 rounds. Post-alignment cleaning consisted of using trimAl (v1.4.rev15) to remove columns containing more than 99% of gaps and manual inspection. The phylogenetic tree was created using IQ-Tree 2 (v2.1.4) with a model of substitution identified using ModelFinder (GTR+F+R10), and optimized trees with nearest neighbor interchange to minimize model violations. Branch support was evaluated with 1000 replicates of SH-aLRT, aBayes, and ultrafast bootstrap support from the IQTREE package. The tree with the highest maximum-likelihood was used as the reconstruction of the group I intron phylogeny. Neighborhood analysis was performed similarly to how the Arc-like ORFs were analyzed. Annotation of rRNA context was extracted from GenBank annotations.

### Generating structures of group I intron RNA structures

The identification of typical group I intron domains was achieved through sequence alignment to the previously characterized *Cdi*IStron (CdISt1) (*31*). Predicted secondary RNA structures were generated using the Mfold (*81*) web server and visualized with the use of RNA canvas (*82*).

### Culturing of *Clostridia senegalense*

A *Clostridia* strain encoding IStrons with ∼80% similarity to the *Cbo*IStron experimentally characterized in this study was obtained from ATCC (strain 25772), where it was defined as belonging to an unknown species classification. Internal rRNA phylogenetic analysis led to the assignment of this strain as a member of species *senegalense*. *C. senegalense* was cultured from a lyophilized ATCC pellet in 5 mL of Gifu Anaerobic Medium Broth, Modified (mGAM; HyServe) in an anaerobic chamber (5% H_2_, 10% CO_2_ and 85% N_2_). All media was pre-reduced for ∼24 hours before use in culturing. *C. senegalense* was then banked as a glycerol stock (final concentration 20%) and sub-cultured into 100 mL cultures of mGAM. The growth of these cultures was monitored with a spectrophotometer over ∼6 h until a final OD_600_ of 0.4-0.6 was reached (exponential phase), at which point cultures were poured into two 50 mL conical tubes and cooled on ice for 10 min. The cultures were then centrifuged at 4,000 *g* for 10 min at 4 °C, the supernatant was decanted, and cell pellets were flash frozen in liquid nitrogen. Pellets were stored at −80 °C until RNA extraction and processing.

### RNA extraction

RNA from *C. senegalense* cell pellets were extracted in 96-well format using a silica bead beating-based protocol adapted from a prior study (*83*). Briefly, 200 µL of 0.1 mm zirconia silica beads (Biospec,) were added to each well of 96-well deep-well plates (Thermo Fisher Scientific). Next, cell pellets were resuspended in 500 µL DNA/RNA shield buffer (Zymo Research) and transferred to separate wells, and the plates were affixed with a sealing mat and centrifuged for 1 min at 4,500 *g*. To avoid overheating, the plates were vortexed for 5 s and incubated at −20 °C for 10 min before bead beating. Then, plates were fixed on a bead beater (Biospec) and subjected to bead beating for 5 min, followed by a 10 min cooling period. The bead beating cycle was repeated three times, and plates were then centrifuged at 4,500 *g* for 5 min to remove cell debris. Next, 60% of the bead beating volume was transferred to the *Quick*-RNA Miniprep Plus kit (Zymo Research), and RNA was purified using the manufacturer’s protocol for gram positive bacteria. RNA quality was assessed using the 260/280 nm ratio (∼2.0) as measured by Nanodrop, and concentration was measured by the Qubit RNA High Sensitivity Assay Kit (Thermo Fisher Scientific) using the manufacturer’s protocol. RNA was stored at −80 °C until library preparation.

### Total RNA sequencing

For total RNA-seq library preparation, 10 µg of purified RNA was treated with Turbo DNase I (Thermo Fisher Scientific) for 1 h at 37 °C using the manufacturer’s protocol. A 2× volume of Mag-Bind TotalPure NGS magnetic beads (Omega) was added to each sample, and the RNA was purified using the manufacturer’s protocol. The RNA was then diluted in NEBuffer2 (NEB) and fragmented by incubating at 92 °C for 1.5 min. To generate RNA with 5’-monophosphate and 3’-hydroxyl ends, samples were treated with RppH (NEB) supplemented with SUPERase•In RNase Inhibitor (Thermo Fisher Scientific) for 30 min at 37 °C, followed by T4 PNK (NEB) in 1× T4 DNA ligase buffer (NEB) for 30 min at 37 °C. Samples were column-purified using RNA Clean & Concentrator-5 (Zymo Research), and the concentration was determined using the DeNovix RNA Assay (DeNovix). Illumina adapter ligation and cDNA synthesis were performed using the NEBNext Small RNA Library Prep kit. Dual index barcodes were added by PCR amplification (12 cycles), and the cDNA libraries were purified using the Monarch PCR & DNA Cleanup Kit (NEB). High-throughput sequencing was performed on an Illumina NextSeq 550 in paired-end mode with 150 cycles per end.

### Whole genome sequencing of *Clostridia senegalense*

Genomic DNA from *C. senegalense* was extracted using the Promega Wizard Genomic DNA purification kit, following the manufacturer’s protocol for gram-positive bacteria. DNA was measured by fluorescent quantification. TnY, a homolog of Tn5, was purified in-house following previous methods (*84*). 10 ng of purified gDNA was tagmented with TnY preloaded with Nextera Read 1 and Read 2 oligos (**Table S4**) following previous methods (*84*), followed by proteinase K treatment (NEB, final concentration 16 U/mL) and column purification. PCR amplification and Illumina barcoding was done for 13 cycles with KAPA HiFi Hotstart ReadyMix (**Table S4**) with an annealing temperature of 63 °C and an extension time of 1 min. The PCR reaction was then resolved on a gel, and a smear from 400–800 bp was extracted for sequencing on a paired end, 150×150 NextSeq kit. Downstream analysis was performed as described in total RNA sequencing. De novo genome assembly was also performed by Plasmidsaurus, and the assembled genome was in agreement with the 4 Mb genome provided for ATCC strain 25772.

### Targeted tagmentation-based detection of IS excision events

100 ng of purified gDNA of *C. senegalense* was tagmented with TnY preloaded with full-length Nextera Read 2/Indexed oligos (**Table S4**). An initial PCR amplification was done with a forward oligo that anneals in the upstream genomic sequence flanking the IStron and an oligo that anneals to the P7 sequence (**Table S4**) using KAPA HiFi Hotstart, with an annealing temperature of 55 °C and 1 min extension time. After bead cleanup using Omega Mag-Bind TotalPure magnetic beads (Omega Bio-Tek) at a ratio of 0.9×, a second PCR was done with an oligo that annealed to the initial PCR amplicon within ∼40 bp of the genome-IStron junction; this forward oligo had all necessary sequences for Illumina sequencing (**Table S4**). After 15 cycles of PCR under the same conditions, the reaction was resolved on a gel, and a smear from 350–800 bp was extracted for sequencing with at least 75 Read 1 cycles. After adapter trimming, the relative abundance of reads that contain a 20-bp sequence of the IStron end a 20-bp sequence of the downstream genomic sequence were tallied using BBDuk from the BBTools suite (v.38.00; https://sourceforge.net/projects/bbmap), with a Hamming distance of 2 and an average Q-score greater than 20.

### RNA-sequencing analyses

RNA-seq datasets were either generated as described above (*C. senegalense* RNA-seq) or downloaded from NCBI (*C. difficile* total RNA-seq: SRR14415999; *C. difficile* small RNA-seq: SRR12329944). RNA-seq data were processed using cutadapt (*85*) (v4.2) to remove adapter sequences, trim low-quality ends from reads, and exclude reads shorter than 18 bp. Reads were mapped to the reference genome (*Cdi*: NZ_CP010905.2; *Cse*: ATCC 25772) using the splice-aware aligner STAR (*86*) (v2.7.10), with --outFilterMultimapNmax 10. Mapped reads were sorted and indexed using SAMtools (*87*) v1.17. Splice junctions inferred by STAR flanking loci of interest were used to create a custom genome annotation file for a second round of STAR alignment, in order to refine splice junction quantitation. Sashimi plots showing read coverage and splice events at specific loci were generated with ggsashimi (*88*) (v1.1.5) in strand-specific mode. Additional read coverage plots were generated by converting alignment files to bigwig files with bamCoverage (*89*) (v3.5.1), using a bin size of 1 and extending reads to fragment size. Coverage over selected genomic regions was visualized in IGV (*90*). For analysis of 3′ boundaries of read pairs, uniquely mapping reads were filtered using SAMtools to only include read 2, and mapping coordinates were extracted using the bamtobed utility from bedtools (*91*) (v2.31.0). The 3′ boundary of each read pair was determined as the start coordinate of read 2, for transcripts on the ‘–’ strand, or the end coordinate of read 2, for transcripts on the ‘+’ strand. To quantify splicing activity at each intron locus, reads were mapped to a mock reference sequence spanning either the 5′ exon1-intron junction, intron-exon2 junction, or the exon-exon junction, using bwa-mem2 (*92*) (v2.2.1). Reads mapping uniquely to each reference sequence were quantified using featureCounts (*93*) (v2.0.2), with a minimum overlap of 3 bp on either end of the junction. Splicing activity was calculated following equation: Splicing activity (%) = N(reads, exon1-exon2)/(N(reads, exon1-exon2)+N_Av_(reads, exon-intron))*100%, where N_Av_ = (N(reads, exon1-intron)+N(reads, intron-exon2))/2.

### RIP-seq

*E. coli* str. K-12 substr. MG1655 (sSL0810) was transformed with plasmids encoding ωRNA and 3×FLAG-CboTnpB (pSL5412) or 3×FLAG-CboTnpB(D189A) (pSL5413). Single colonies were inoculated in liquid LB with spectinomycin (100 µg mL^−1^) and grown overnight. The next day, the culture was used to inoculated 50 mL of liquid LB with spectinomycin (100 µg mL^−1^) at 100× dilution and grown until OD_600_ reached 0.5. 10 mL of culture was centrifuged at 4,000 *g* for 10 min at 4 °C, and the supernatant was removed. The pellet was washed once with 1 mL of cold TBS (20 mM Tris-HCl (pH 7.5 at 25 °C), 150 mM NaCl), centrifuged at 10,000 *g* for 5 min at 4 °C, the supernatant was removed, and the resulting pellet was flash-frozen in liquid nitrogen. Pellets were stored at −80 °C. Antibodies for immunoprecipitation were conjugated to magnetic beads as follows: for each sample, 30 μL Dynabeads Protein G (Thermo Fisher Scientific) were washed 3× in 1 mL RIP lysis buffer (20 mM Tris-HCl (pH 7.5 at 25 °C), 150 mM KCl, 1 mM MgCl_2_, 0.2% Triton X-100), resuspended in 1 mL RIP lysis buffer, combined with 10 μL anti-FLAG M2 antibody (Sigma-Aldrich), and rotated for >3 hours at 4 °C. Antibody-bead complexes were washed 3× to remove unconjugated antibodies, and resuspended in 30 μL RIP lysis buffer per sample.

To generate cell lysates, flash-frozen pellets were first resuspended in 1.2 mL RIP lysis buffer supplemented with cOmplete Protease Inhibitor Cocktail (Roche) and SUPERase•In RNase Inhibitor (Thermo Fisher Scientific). Cells were then sonicated for 1.5 min total (2 s ON, 5 s OFF) at 20% amplitude. To clear cell debris and insoluble material, lysates were centrifuged for 15 min at 4 °C at 21,000 *g*, and the supernatant was transferred to a new tube. At this point, a small volume of each sample (24 μL, or 2%) was set aside as the “input” starting material and stored at −80 °C.

For immunoprecipitation, each sample was combined with 30 μL antibody-bead complex and rotated overnight at 4 °C. The next day, each sample was washed 3× with ice-cold RIP wash buffer (20 mM Tris-HCl (pH 7.5 at 25 °C), 150 mM KCl, 1 mM MgCl_2_). After the last wash, beads were resuspended in 1 mL TRIzol (Thermo Fisher Scientific) and incubated at RT for 5 min to allow separation of RNA from the beads. A magnetic rack was used to isolate the supernatant, which was transferred to a new tube and combined with 200 μL chloroform. Each sample was mixed vigorously by inversion, incubated at RT for 3 min, and centrifuged for 15 min at 4 °C at 12,000 *g*. RNA was isolated from the upper aqueous phase using the RNA Clean & Concentrator-5 kit (Zymo Research) and eluted in 15 μL RNase-free water. RNA from input samples was isolated in the same manner using TRIzol and column purification.

For RIP-seq library preparation (input and RIP eluates), 6 μL of RNA was diluted in FastAP Buffer (Thermo Fisher Scientific) supplemented with SUPERase•In RNase Inhibitor (Thermo Fisher Scientific) to a total volume of 18 μL, and fragmented by heating to 92 °C for 1.5 min. Each sample was treated with 2 μL TURBO DNase (Thermo Fisher Scientific) for 30 min at 37 °C and column-purified using the RNA Clean & Concentrator-5 kit (Zymo Research), eluting in 12.5 μL RNase-free water. RNA concentration was quantified using the DeNovix RNA Assay. Illumina sequencing libraries were prepared using the NEBNext Small RNA Library Prep kit, and libraries were sequenced on an Illumina NextSeq 500 in paired-end mode with 75 cycles per end.

Adapter trimming, quality trimming, and read length filtering of RIP-seq reads was performed as described above for total RNA-seq experiments. Trimmed and filtered reads were mapped to a reference containing both the MG1655 genome (NC_000913.3) and plasmid (pSL5412) sequences using bwa-mem2 (*94*) (v2.2.1) with default parameters. SAMtools (v1.17) was used to filter uniquely mapped reads (MAPQ > 1), as well as used to sort and index the uniquely mapped reads. Coverage tracks were generated using bamCoverage (v3.5.1) with a bin size of 1, read extension to fragment size, and normalization by counts per million mapped reads (CPM) with exact scaling. Coverage was visualized in IGV.

### Sequence identity matrices

Pairwise sequence identity matrices were generated in Geneious from MAFFT alignments of intron or ωRNA nucleic acid sequences, or TnpA or TnpB amino acid sequences, using default settings. For introns, the identity matrices used the conserved structured portion of each IStron, determined by multiple sequence alignment, and the Group I intron boundaries determined by splicing analysis of RNA-seq data or predicted by the intron covariance model. For ωRNAs, the identity matrices used the ωRNA boundaries determined by RNA-seq (for *C. senegalense*), or the 117 nt downstream of TnpB (for *C. difficile*, due to poor resolution of ωRNA boundaries in the public RNA-seq datasets analyzed in this study). Accession numbers for IStron, TnpA, and TnpB sequences are listed in **Table S1.**

### Plasmid and *E. coli* strain construction

All strains and plasmids used in this study are described in **Tables S2 and S3**, respectively, and a subset is available from Addgene. Briefly, genes encoding *Cbo*TnpA and native *Cbo*IStron sequence were synthesized by Twist Bioscience. *E. coli* codon-optimized *Cbo*TnpB and bioinformatically predicted ωRNA were synthesized and cloned into a single pCDF-Duet vector by Genscript, with two separate J-23 series promoters driving their expression. Transposase expression plasmids were generated using Gibson assembly, by inserting the *tnpA* gene downstream of pLac or T7 promoters in a minimal pCOLADuet-1 vector (*7*). Native IStron, IStron with TnpB only and mini-IS sequence (581 bp from the left end and 221 bp of the right end) were cloned using Gibson assembly, by inserting them into a pCDF-Duet vector downstream of T7 promoter. pTarget plasmids were generated by around-the-horn PCR, inserting a 44-bp target sequence into a minimal pCOLA-Duet vector. A transposition circular intermediate (pDonor_CI_) was generated by Gibson assembly of the *Cbo*IStron left end (581 bp), right end (221 bp), R6K ori, and chloramphenicol resistance gene. The cloning mix was used to transform a *pir+* strain (sSL0281, CGSC), to allow for the propagation of the R6K ori-bearing plasmid. Derivatives of these plasmids were cloned using a combination of methods, including Gibson assembly, restriction digestion-ligation, ligation of hybridized oligonucleotides, Golden Gate Assembly, and around-the-horn PCR. Plasmids were cloned and propagated in NEB Turbo cells (NEB) — except for pDonor_CI_ derivatives, which were propagated in strain sSL0281 — purified using Miniprep Kit (Qiagen), and verified by Sanger sequencing (GENEWIZ).

### DNA cleavage assays with TnpB

Plasmid interference assays were performed in *E. coli* str. K-12 substr. MG1655 (sSL0810) when a synthetic *Cbo*TnpB expression construct was used, and in *E. coli* BL21(DE3) strain for all other experiments. When *Cbo*TnpB was co-expressed with ωRNA from the same plasmid, BL21(DE3) cells were transformed with a pEffector plasmid, and single colony isolates were selected to prepare chemically competent cells. 200 ng of pTarget plasmid were then delivered via transformation. After 2 h, cells were spun down at 6,000 rpm for 5 min and resuspended in 30 µL of LB. Cells were then serially diluted (10×) and plated on LB-agar media containing spectinomycin (100 µg mL^−1^) and kanamycin (50 µg mL^−1^), and grown for 24 h at 37 °C. Plates were imaged in an Amersham Imager 600. For experiments when a mini-IS was used as a guide for *Cbo*TnpB, BL21(DE3) cells were co-transformed with mini-IS and TnpB expression plasmids, and single colony isolates were selected to prepare chemically competent cells. A second transformation was performed as indicated previously, and cells were plated on LB-agar media containing spectinomycin (100 µg mL^−1^), chloramphenicol (25 µg mL^−1^), kanamycin (50 µg mL^−1^) and IPTG (0.1 mM), and grown for 24 h at 37 °C. Plates were imaged in an Amersham Imager 600.

### TAM library experiments and analyses

TAM library experiments were prepared for sequencing, as previously described (*7*). Analyses were also performed as previously described. In brief, reads were filtered on containing the correct sequence both upstream and downstream of the TAM region. TAM sequences were then extracted, tallied, and depletion values were calculated as the relative abundance of the library member in the output library divided by the relative abundance of the library member in the input. Sequence logos were generated with the library members that were depleted more than 5-fold (depletion value greater than 32) using WebLogo (*95*) (v2.8), and the top 5% of depleted library members were used to generate TAM wheels (depletion scores were inverted to properly visualize TAM wheels) (*96*).

### Transposon excision assays with TnpA_S_

All transposition experiments were performed in *E. coli* MG1655 strains. Chemically competent cells were first transformed with a plasmid encoding *tnpA* under an inducible *lac* promoter, and transformants were isolated by plating on LB-agar plates with antibiotic (50 µg mL^−1^ kanamycin). Liquid cultures were then inoculated from single colonies, and the resulting strains were made chemically competent using standard methods, aliquoted, and snap-frozen. This strain was then subsequently transformed with 100 ng of a separate pDonor plasmid containing a Mini-IS variant. Cultures were grown overnight at 37 °C on LB-agar plates under antibiotic selection (100 µg mL^−1^ spectinomycin, 50 µg mL^−1^ kanamycin) and IPTG induction (0.5 mM). The next day, all colonies were scraped and resuspended in LB medium. To prepare cell lysates, approximately 3.2 × 10^8^ cells (equivalent to 200 μL of culture at OD_600_ = 2.0) were transferred to a 96-well plate. Cells were pelleted by centrifugation at 4,000 *g* for 5 min and resuspended in 80 μL of H_2_O. Next, cells were lysed by incubating at 95 °C for 10 min in a thermal cycler. The cell debris was pelleted by centrifugation at 4,000 *g* for 5 min, and 10 μL of lysate supernatant were removed and serially diluted in H_2_O to generate 10- and 100-fold lysate dilutions for PCR and qPCR analyses, respectively.

PCR reactions were performed using primers annealing in 5′ and 3′ IStron-flanking sequences, which can be used to amplify both unexcised loci (longer amplicon) and excised loci (shorter amplicon). Each 20 µL PCR reaction contained 1× OneTaq Master Mix (NEB), 0.2 µM of each primer, and 1 µL of 10-fold diluted lysate. Thermal cycling conditions included DNA denaturation (94 °C for 30 s), 30 cycles of amplification (denaturation: 95 °C for 15 s, annealing: 46 °C for 15 s, extension: 68 °C for 15 s), followed by a final extension (68 °C for 5 min). Products were resolved by 1.5% agarose gel electrophoresis and visualized by staining with SYBR Safe (Thermo Fisher Scientific). Quantitative PCR (qPCR) was performed in a 10 µL reaction that contained 5 µL SsoAdvanced™ Universal SYBR Green Supermix (BioRad), 1 µL H_2_O, 2 µL of primer pair at 2.5 µM concentration, and 2 µL of 100-fold diluted lysate. Two primer pairs were used: (1) excision products were specifically captured using a forward primer annealing to 5′ IStron flanking sequence (oSL12715) and reverse primer spanning the donor joint (oSL12719); (2) a reference gene (*specR*) was amplified with primers annealing to the same pDonor plasmid (oSL13566 and oSL13567). Excision efficiency (%) was calculated as a ratio between experimental sample and a lysate from cells transformed with a mock plasmid (mimicking donor joint), using the formula: excision efficiency (%) = 2^-ΔΔCq^*100%, where ΔΔCq = ΔCq(Cq_sample_(donor joint)-(reference gene))-ΔCq(Cq_mock_(donor joint)-(reference gene)). Primer sequences are provided in **Table S4**.

### TnpA_S_ transposition intermediate (minicircle) capture using PCR

A transposition assay was performed in *E. coli* BL21(DE3) strain transformed with pDonor plasmid encoding mini-IS with *cmR* gene as a cargo (pSL5192). Cells were made chemically competent and transformed with plasmids encoding *tnpA* (pSL5006), mutant *tnpA* (pSL5094), or an empty vector (pSL4032). Transformants were plated on LB-agar plates with antibiotic (100 µg mL^−1^ carbenicillin, 50 µg mL^−1^ kanamycin). The next day, several colonies were used to inoculate liquid cultures in LB with 100 µg mL^−1^ carbenicillin, 50 µg mL^−1^ kanamycin, and grown overnight. An aliquot of turbid culture equivalent of 200 µL of culture at OD_600_ = 2.0 was taken for lysis. Lysis was performed as described previously, and minicircle junction was amplified using a forward primer annealing on transposon RE, and a reverse primer annealing on the transposon LE (see **Table S4** for primer sequences). 20 µL PCR reaction contained 1× OneTaq Master Mix (NEB), 0.2 µM of each primer, and 1 µL of 10-fold diluted lysate. Thermal cycling: DNA denaturation (94 °C for 30 s), 30 cycles of amplification (denaturation: 95 °C for 15 s, annealing: 51 °C for 15 s, extension: 68 °C for 20 s), followed by a final extension (68 °C for 5 min). Products were resolved by 1.5% agarose gel electrophoresis and visualized by staining with SYBR Safe (Thermo Fisher Scientific). The expected junction was confirmed by Sanger sequencing.

### Transposon integration assays with TnpA_S_

Plasmids with an R6K ori, a *CmR* marker, and inverted IStron ends (pDonor_CI_) were cloned in *pir+* strain (sSL0281). *E. coli* str. K-12 substr. MG1655 transformed with either a TnpA or mutant TnpA expression plasmid driven by pLac was grown in liquid LB with IPTG (0.5 mM), and cells were first made chemically competent. These cells were transformed with a pDonor_CI_ with a canonical GG core dinucleotide. After recovery for 7 h, cells were plated on LB-agar plates with chloramphenicol (25 µg mL^−1^) and grown for ∼24 hours. The number of surviving colonies were counted across replicates. Surviving colonies from WT TnpA-expressing cells were then scraped for downstream sequencing. To investigate the impact of core dinucleotide modifications, MG1655 cells with a WT TnpA expression plasmid driven by pLac were grown in liquid LB with IPTG (0.5 mM) and made electrocompetent at OD_600_ = 0.6. These cells were then electroporated with pDonor_CI_ variants, recovered for 7 h, plated on LB-agar plates with chloramphenicol (25 µg mL^−1^), and grown for ∼24 hours. Surviving colonies were pooled, and genomic DNA was extracted and quantified via Qubit. Approximately 100 ng of gDNA was tagmented with TnY pre-loaded with Read 2 Nextera oligos (**Table S4**). Two rounds of PCR were performed as described for targeted detection of IS excision events with oligos that annealed to either the IStron left end or right end (**Table S4**). Paired-end, 76×76 cycle sequencing was performed on a NextSeq platform. Using BBDuk (*97*), reads were then filtered for containing the proper IStron end sequence, and the flanking genomic sequence was extracted. Reads that contained the parental pDonor_CI_ sequence were removed during this process. Flanking genomic sequences were then aligned to the *E. coli* genome using Bowtie2 (*98*). WebLogo (*95*) representations were generated using the input sequence on both the IStron left end and right end, as well as the mapped genomic insertion sites.

### Transposon maintenance experiments with TnpA_S_ and TnpB

sSL3391, a derivative of *E. coli* str. K-12 substr. MG1655 with *lacZ* deletion replaced by a chloramphenicol resistance cassette, was transformed with 400 ng of plasmid encoding an intact *lacZ* gene (pSL4825, empty vector) or a CboIStron-interrupted *lacZ* gene (pSL5948, pSL5949, pSL5950; see **Table S3** for descriptions). Following transformation, colonies were plated on MacConkey agar media containing tetracycline (10 µg mL^−1^) to enrich for IStron excision events. Cells were grown at 37 °C for 36 h, then harvested, serially diluted, plated onto LB-agar containing tetracycline (10 µg mL^−1^) and X-gal (200 μg mL^−1^), and grown for 18 h at 37 °C. The total number of colonies were counted, along with the number of blue colonies to determine the frequency of excision and reintegration events. Colony counts are represented as colony forming units (CFUs) per μg of DNA. In addition, genomic lysates were harvested from cells, as described in the assay for excision detection, for PCR analysis.

### Transposon recombination assay with TnpA_S_ and TnpB

*E. coli* str. K-12 substr. MG1655 (sSL0810) containing an intact *lacZ* loci were chemically transformed with 400 ng of plasmid encoding an intact *lacZ* gene (pSL4825, empty vector) or *Cbo*IStron-interrupted *lacZ* gene (pSL5948, pSL5949, pSL5950; see **Table S3** for descriptions), recovered for 1 h at 37 °C in liquid LB, and serially diluted on LB-agar plates with tetracycline (10 µg mL^−1^). The next day, colonies were counted, and the tetracycline plates were replica plated to LB-agar plates containing both tetracycline (10 μg mL^−1^) and X-gal (200 μg mL^−1^) for blue/white colony screening. White colonies were counted to determine the frequency of recombination events at the genomic *lacZ* locus. All colony counts are represented as colony forming units (CFUs) per μg of DNA.

### *In vitro* splicing assays

Templates for *in vitro* splicing reactions were obtained by PCR amplification of mock excised (pSL5516), splicing mutant (pSL5026), and mini-IS (pSL5515) containing plasmids. All templates had a T7 promoter encoded within the amplicon, which is required for transcription (see **Table S4** for primer sequences). PCR products were extracted after gel electrophoresis, and 1 µg of each construct was used in 50 µL *in vitro* transcription reaction. Reactions comprised 30 mM Tris (pH 8.0 at 25 °C), 10 mM DTT, 0.1% Triton X-100, 0.1% spermidine, 60 mM MgCl_2_, 0.2 µL SUPERase•In™ (Thermo Fisher Scientific), 6 mM each NTP and 0.2 mg/mL of T7 polymerase. Reactions were incubated overnight at 37 °C. The next day, pyrophosphate precipitate was removed by centrifugation, and the DNA template was digested by adding 1 µl of TURBO™ DNase (2 U/µL) (Thermo Fisher Scientific) and incubating for 30 min at 37 °C. The resulting RNA was purified using the Monarch RNA Cleanup Kit (NEB) and stored at −80 °C.

### *In vivo* splicing assays

*In vivo* splicing assays were performed in *E. coli* BL21 (DE3) strain transformed with a mini-IS variant encoding plasmid, or co-transformed with mini-IS and TnpB expression plasmids. For single-plasmid transformations, single colonies were picked from a plate and used to inoculate overnight cultures in LB with spectinomycin (100 µg mL^−1^). In the morning, the cultures were re-inoculated at 40× dilution in LB supplemented with spectinomycin (100 µg mL^−1^) and IPTG (0.1 mM), and grown until the OD_600_ reached 0.5-0.7. Then, an aliquot equivalent to 250 µL of cell suspension at OD_600_ = 0.5 was taken from each culture and centrifuged at 6,000 rpm for 5 min, and the cell pellet was resuspended in 750 µL Trizol (Thermo Fisher Scientific). After incubating 10 min at room temperature, 150 µL of chloroform was added, and tubes shaken and centrifuged at 12,000 *g* for 15 min at 4 °C. The aqueous phase was transferred to a new tube and mixed with an equal volume of absolute ethanol (>96%), followed by RNA purification using the Monarch RNA Cleanup Kit (NEB). Purified RNA was stored at −80 °C. For splicing assays with TnpB co-expressed *in trans*, single colonies were used to inoculate overnight cultures in LB with spectinomycin (100 µg mL^−1^) and chloramphenicol (25 µg mL^−1^). In the morning, the cultures were re-inoculated at 40× dilution in LB supplemented with spectinomycin (100 µg mL^−1^), chloramphenicol (25 µg mL^−1^) and IPTG (0.5 mM), and grown until OD_600_ reached 0.5-0.7. All downstream steps were performed as described above.

### Reverse transcription (RT)-PCR and RT-qPCR analyses

200 ng of the purified total RNA was used as an input for reverse transcription reactions. First, total RNA was treated with 1 µL of dsDNase (Thermo Fisher Scientific) in 1× dsDNase reaction buffer in the final volume of 10 µL, incubating at 37 °C for 20 min. Then 1 µL of 10 mM dNTP, 1 µL of 2 µM oSL12027, and 1 µl of 2 µM oSL13568 were added for gene-specific priming, and samples were heated at 65 °C for 5 min. Tubes were then placed directly on ice, followed by addition of 4 µL of SSIV buffer, 1 µL 100 mM DTT, 1 µL SUPERase•In™ (Thermo Fisher Scientific) and 1 µL of SuperScript IV Reverse Transcriptase (200 U/µL, Thermo Fisher Scientific), incubation at 53 °C for 10 min, and then incubation at 80 °C for 10 min. The resulting cDNA was diluted and used for end-point or quantitative PCR.

Endpoint PCR was performed in a 20 µL reaction volume containing 1× OneTaq Master Mix (NEB), 0.2 µM of each primer and 1 µL of 100-fold diluted cDNA. Thermal cycling conditions were as follows: DNA denaturation (94 °C for 30 s), 30 cycles of amplification (denaturation: 95 °C for 15 s, annealing: 46 °C for 15 s, extension: 68 °C for 15 s), followed by a final extension (68 °C for 5 min). Products were resolved by 1.5% agarose gel electrophoresis and visualized by staining with SYBR Safe (Thermo Fisher Scientific). We verified that splicing was not a result of residual DNA excision products being formed in cells (e.g. from trans-acting *E. coli* transposases) by carefully verifying the absence of any junction products under ΔTnpA_S_ conditions in transposon excision assays (**Fig. 2B**).

Quantitative PCR (qPCR) was performed in 10 µL reactions containing 5 µL SsoAdvanced™ Universal SYBR Green Supermix (BioRad), 1 µL H_2_O, 2 µL of primer pair at 2.5 µM concentration, and 2 µL of 100-fold diluted cDNA (10-fold when intron was expressed from a J23114 promoter). Two primer pairs were used: (1) spliced RNAs were captured using a forward primer annealing to exon 1 (oSL12715) and reverse primer spanning the splice-junction (oSL12719); (2) unspliced products were amplified using the same forward primer annealing to exon 1 (oSL12715) and reverse primer annealing to the IStron left end (oSL13922). Reactions were prepared in 384-well clear/white PCR plates (BioRad), and measurements were performed on a CFX384 RealTime PCR Detection System (BioRad) using the following thermal cycling parameters: polymerase activation and DNA denaturation (98 °C for 2.5 min), 40 cycles of amplification (98 °C for 10 s, 62 °C for 20 s), and terminal melt-curve analysis (decrease from 95 °C to 65 °C in 0.5 °C/5 s increments). For each sample, the ratio of spliced/unspliced was obtained by calculating spliced/unspliced = 2^-ΔCq^, where ΔCq = Cq(spliced)-Cq(unspliced). All primer sequences are provided in **Table S4**.

## ACKNOWLEDGEMENTS

We thank Z. Akhtar for laboratory support, N. Sanjana for help with TnY-based tagmentation assays, A.-L. Steckelberg for help with *in vitro* transcription reactions, A. Bernheim and C. Le Maréchal for helpful discussions about *C. botulinum*, H.H. Wang for anaerobic chamber access for *C. senegalense* culturing, L.F. Landweber for qPCR instrument access, and the JP Sulzberger Columbia Genome Center for NGS support.

## Author contributions

R.Z., C.M., and S.H.S. conceived of and designed the project. R.Z. developed excision assays and performed all TnpB and splicing assays. H.C.L. performed most bioinformatics analyses, with support from C.M. and S.T. S.T. performed and analyzed RIP-seq and RNA-seq experiments. C.M. selected experimental IStron systems and performed transposon retention/recombination assays. G.D.L. performed and analyzed tagmentation, TAM library, and dinucleotide swapping experiments. E.E.M. performed and analyzed transposon excision assays and group I intron secondary structures. S.R.P. developed and performed initial splicing assays. D.R.G. cultured *C. senegalense* and prepared RNA samples. T.W. assisted with bioinformatics and figure generation. R.Z., C.M., and S.H.S. discussed the data and wrote the manuscript, with input from all authors.

## Data and materials availability

Next-generation sequencing data are made available in the National Center for Biotechnology Information (NCBI) Sequence Read Archive. Datasets generated and analyzed in the current study are available from the corresponding author upon reasonable request.

## SUPPLEMENTARY MATERIALS

### Materials and Methods

**Fig. S1.**
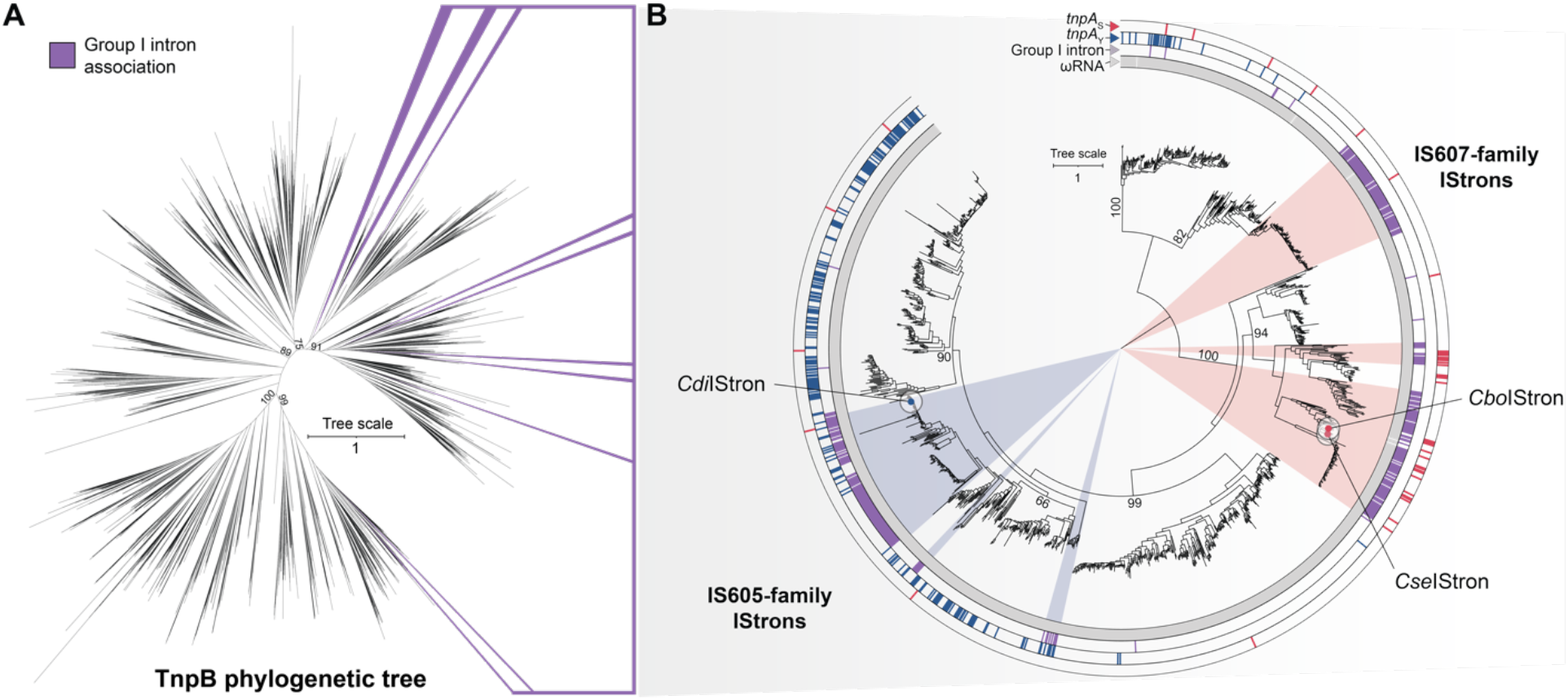
Evolutionary and neighborhood analyses of TnpB, TnpA, and group I introns. (**A**) Unrooted phylogenetic tree of bacterial TnpB homologs from Meers *et al.* (*7*), in which cluster representatives are highlighted (purple) that contain any member associated with a group I intron. Bootstrap values are indicated for major nodes. (**B**) Focused phylogenetic tree of TnpB homologs from **A**, including a much larger set of additional representatives from all clusters. Neighborhood analyses were performed on the genomic contexts of each *tnpB* gene, revealing associations with *tnpA*_S_ (IS607-family), *tnpA*_Y_ (IS200/IS605-family), group I introns (IStron), and ωRNA loci. Large groups of IS607- (red sector) and IS200/IS605-family (blue sector) IStrons were identified, and representative *Cdi*IStron, *Cbo*IStron, and *Cse*IStron members are annotated. Bootstrap values are indicated for major nodes.

**Fig. S2.**
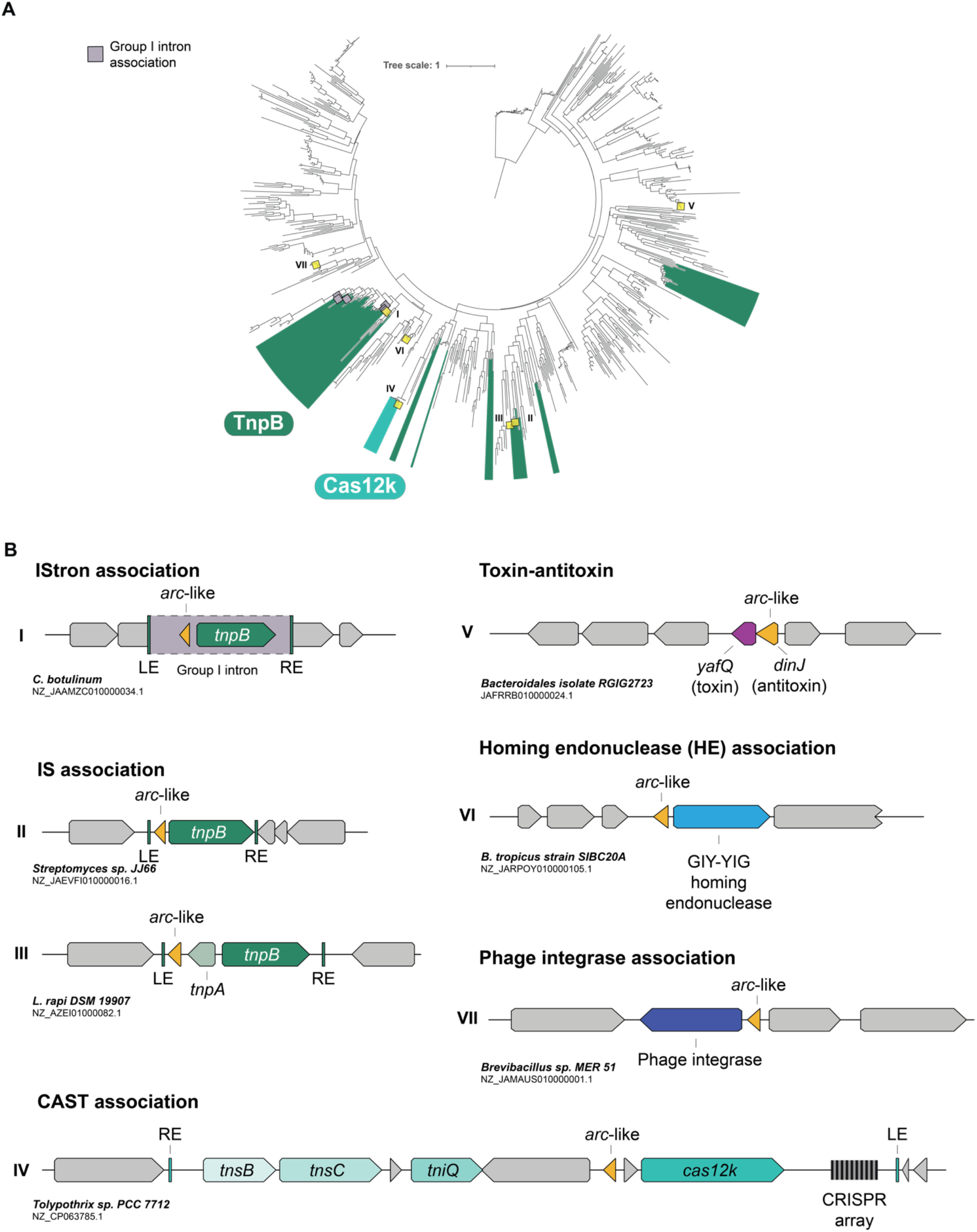
Evolutionary and neighborhood analyses of transposon-associated Arc-like proteins. (**A**) Phylogenetic tree of Arc-like proteins, revealing genetic associations with TnpB (IS-family transposons) and Cas12k (CRISPR-associated transposons) alongside many other classes of mobile genetic elements. (**B**) Genetic architecture of representative transposable elements encoding Arc-like proteins (yellow arrow), including IStrons, ISs, CRISPR-associated transposons (CASTs), toxin-antitoxin (TA) systems, homing endonucleases (HEs), and phage integrases. Relevant genes are annotated, and putative transposon boundaries are indicated with green boxes. Roman numerals next to each representative genomic locus correspond to positions annotated on the phylogenetic tree in **A**.

**Fig. S3.**
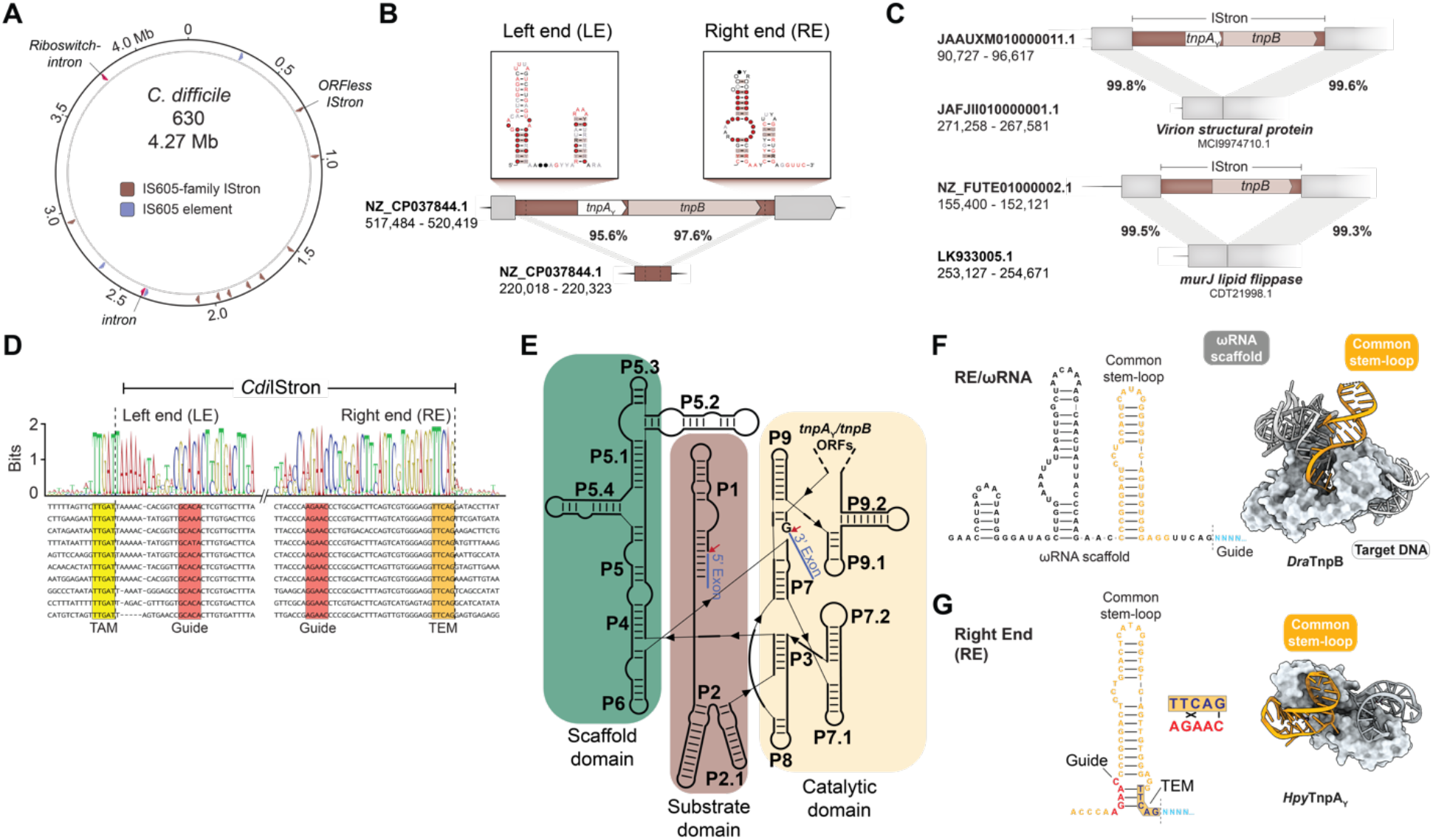
Genomic and functional analysis of IS200/IS605-family IStrons from *C. difficile* (*Cdi*IStron). (**A**) Genome-wide distribution of group I introns and IS200/IS605-family transposons in *C. difficile* strain 630 (NCBI accession ID: NZ_CP010905.2). *Cdi*IStron copies are colored distinctly from standalone group I introns and an IS605-family element. (**B**) Transposon left end (LE) and right end (RE) definitions for a representative *Cdi*IStron element (top), based on covariance models; the predicted LE and RE secondary structures recognized by TnpA_Y_ during transposon excision and integration are shown in the inset (top). A palindrome-associated transposable element (PATE) harboring IStron ends, but no open reading frames, is shown below. (**C**) Comparative genomics analysis of two *Cdi*IStron loci, showing the same location in isogenic strains that do not have the IStron insertion. This analysis corroborates the LE/RE boundary assignments from **B.** (**D**) DNA multiple sequence alignment of transposon left end (LE) and right end (RE) sequences for 10 select *Cdi*IStrons, based on comparative genomics and covariance models, with a consensus sequence shown at the top. The 5′-TTGAT-3′ transposon adjacent motif (TAM), the transposon encoded motif (TEM), and DNA guide sequences for both LE and RE are highlighted in yellow, orange, and red, respectively; dotted black lines indicate the upstream and downstream transposon boundaries. (**E**) Secondary structure of the group I intron from a representative *Cdi*IStron, with scaffold, substrate, and catalytic domains colored in green, brown, and yellow, respectively. Paired stem-loops defined as P1–P9, according to conventions defined by Hasselmayer *et al.* (*31*); the region that harbors *tnpA*_Y_ and/or *tnpB* ORFs is indicated, as are the predicted 5′SS and 3′SS (red arrows); flanking exons are labeled. (**F**) Predicted ωRNA secondary structure for a representative *Cdi*IStron from **A**, based on secondary structure folding and alignment to the covariance model. The region predicted to be important for 3′ SS recognition is highlighted, and the region also recognized by TnpA_Y_ at the DNA level is highlighted in orange. A cryoEM structure of *Dra*TnpB (ISDra2) bound to its ωRNA substrate (PDB ID: 8H1J) (*57*) is shown at right, highlighting the stem-loop (orange) that is recognized similarly at the RNA and DNA levels by TnpB and TnpA_Y_, respectively. (**G**) Secondary structure of the transposon RE ssDNA for the same *Cdi*IStron from **F**, based on covariance modeling; predicted DNA-DNA base-pairing between the DNA ‘guide’ (red) and transposon-encoded motif (TEM; orange) is highlighted. An X-ray crystal structure of *Hpy*TnpA_Y_ bounds to its RE substrate (PDB ID: 2VJU) (*53*) is shown at right, highlighting the stem-loop (orange) that is recognized similarly at the RNA and DNA levels by TnpB and TnpA_Y_, respectively.

**Fig. S4.**
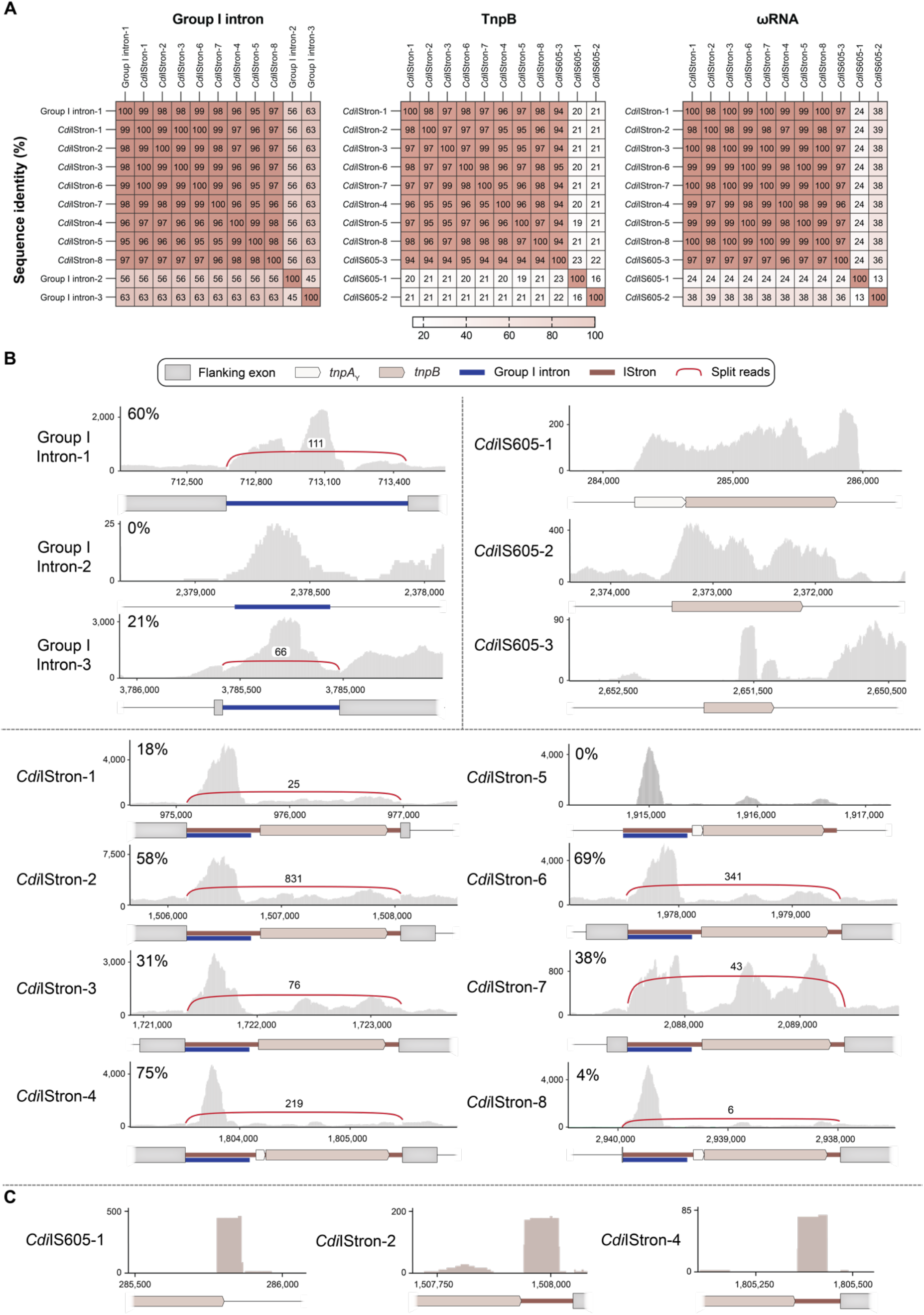
Comparative sequence and RNA-seq analyses of *C. difficile* intron, IStron, and IS605 elements. (**A**) Pairwise sequence identity matrices for all identifiable group I introns (left), TnpB homologs (middle), and ωRNAs (right) encoded by *C. difficile* strain 630 (NCBI accession ID: NZ_CP010905.2), with percent identities shown in the heat maps. Rows and columns were grouped using hierarchical clustering. Group I introns and ωRNAs were compared using nucleotide sequence, TnpB – using amino acid sequence. Three standalone group I introns (denoted 1–3) are not encoded within IS elements, and all *Cdi*IStrons fall within the IS200/IS605 family. Three additional IS605-family transposons which do not encode group I introns are shown. (**B**) Read coverage from a published total RNA-seq dataset at the indicated *Cdi*IStron loci in *C. difficile* strain 630, as well as a representative standalone group I introns (left) and non-intron-containing IS605-family elements (right). The genomic coordinates are shown below each plot, as are gene annotations. Non-uniquely mapping reads are shown, due to the high level of sequence identity between loci that results in loss of coverage when plotting uniquely mapping reads. RNA-seq coverage corresponding to the folded portion of group I introns are indicated (blue). The number and connectivity of spliced exon-exon junction reads are highlighted (red), and quantitative comparison of exon-intron and exon-exon reads yields an apparent splicing percentage at the RNA level (top left of each plot). Splice event quantitation is not affected by high levels of sequence identity between IStron elements, as the flanking exons are unique. These analyses reveal a wide range of *Cdi*IStron splicing efficiencies, though numbers may also be affected by DNA transposon excision events within the population. (**C**) Read coverage from a published small RNA-seq dataset corresponding to a portion of the ωRNA for representative loci are shown. Coverage over the ωRNA was not easily interpretable due to high sequence identity between the IStrons, leading to ambiguous mapping; furthermore, the 3’ boundary of the ωRNA cannot be resolved from this dataset due to the use of single-end sequencing. RNA-seq data in **B** and **C** were previously published (*50*, *51*).

**Fig. S5.**
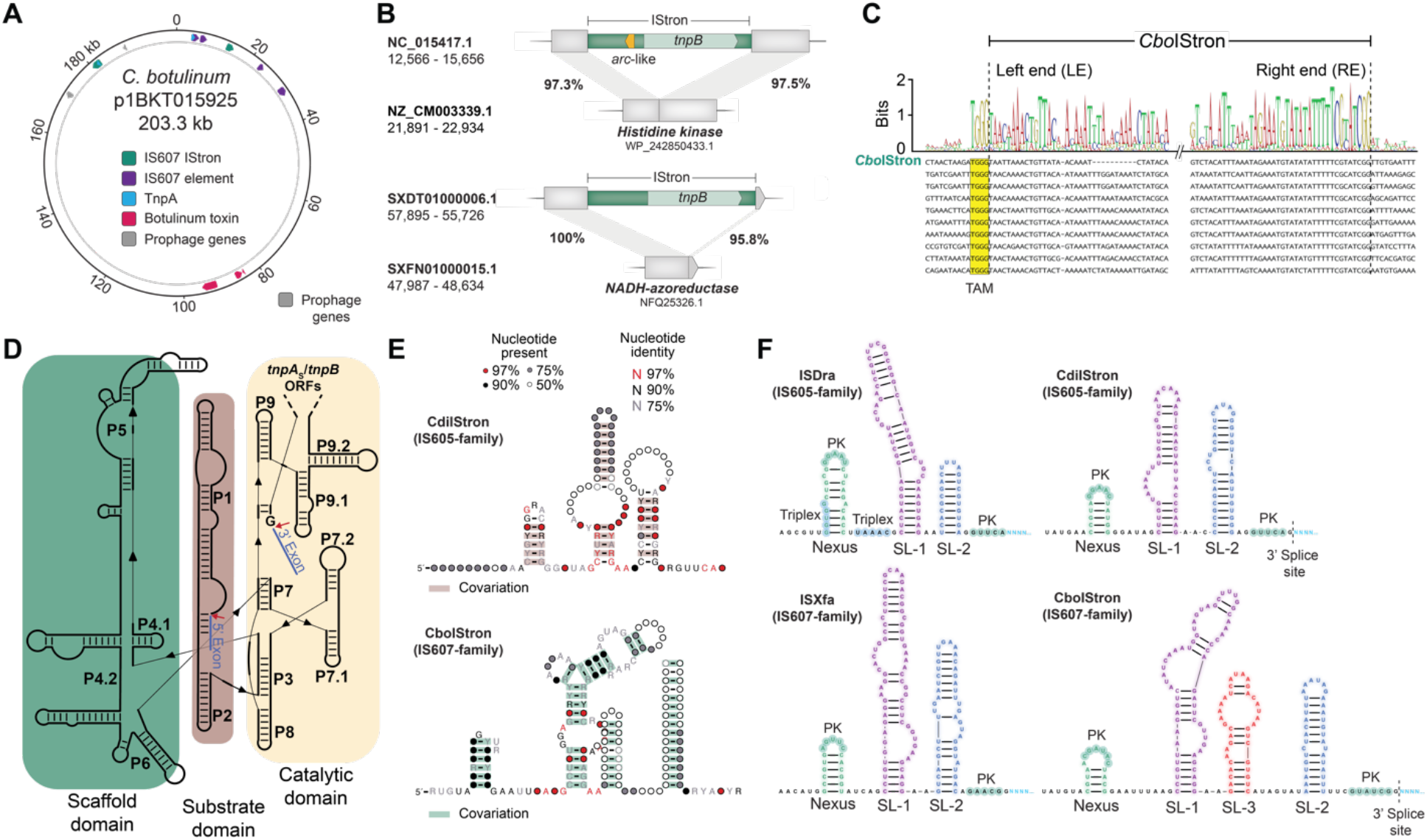
Genomic and functional analysis of IS607-family IStrons from *C. botulinum* (*Cbo*IStron). (**A**) Schematic of a plasmid (NCBI accession ID: NC_015417.1) in *C. botulinum* strain BKT015925, highlighting the location of the botulinum neurotoxin gene and IS605-family, IS607-family, and IS607-family IStron elements. (**B**) Examples of IStron-interrupted protein coding genes in *C. botulinum*, with isogenic strains showing an intact protein coding gene with no IStron insertion. (**C**) DNA multiple sequence alignment of transposon LE and RE sequences for 10 selected *Cbo*IStrons, with a consensus sequence shown at the top. The predicted 5′-TGGG-3′ transposon adjacent motif (TAM) is highlighted in yellow; dotted black lines indicate the upstream and downstream transposon boundaries. The top row corresponds to *Cbo*IStron, which is the source of the TnpA_S_, TnpB, ωRNA, and intron constructs used in heterologous *E. coli* experiments. (**D**) Secondary structure of the group I intron from a representative *Cbo*IStron, with scaffold, substrate, and catalytic domains colored in green, brown, and yellow, respectively. Paired stem-loops defined as P1–P9, according to conventions defined by Hasselmayer *et al.* (*31*); the region that harbors *tnpA_S_* and/or *tnpB* ORFs is indicated, as are the predicted 5′SS and 3′SS (red arrows); flanking exons are labeled. (**E**) Covariance models for the *Cdi*IStron (top) and *Cbo*IStron (bottom) ωRNA. (**F**) Comparison of ωRNAs from representative IS605- and IS607-family transposons from *D. radiodurans* and *Xylella fastidiosa* (left), as well as IS605- and IS607-family IStrons from *C. difficile* and *C. botulinum* (right). Distinct RNA secondary structure motifs are labeled, alongside predicted pseudoknot (PK) interactions, and the beginning of the ωRNA guide sequence at the 3′ end is shown in blue. For IStrons, the guide sequence immediately follows the predicted 3′SS. The nexus SL predicted to form a pseudo-knot with the ωRNA scaffold 3’ end, and other stem loops are numbered from left to right. An exception is made for *Cbo*IStron, which has an additional stem loop between SL1 and SL2, denoted SL3.

**Fig. S6.**
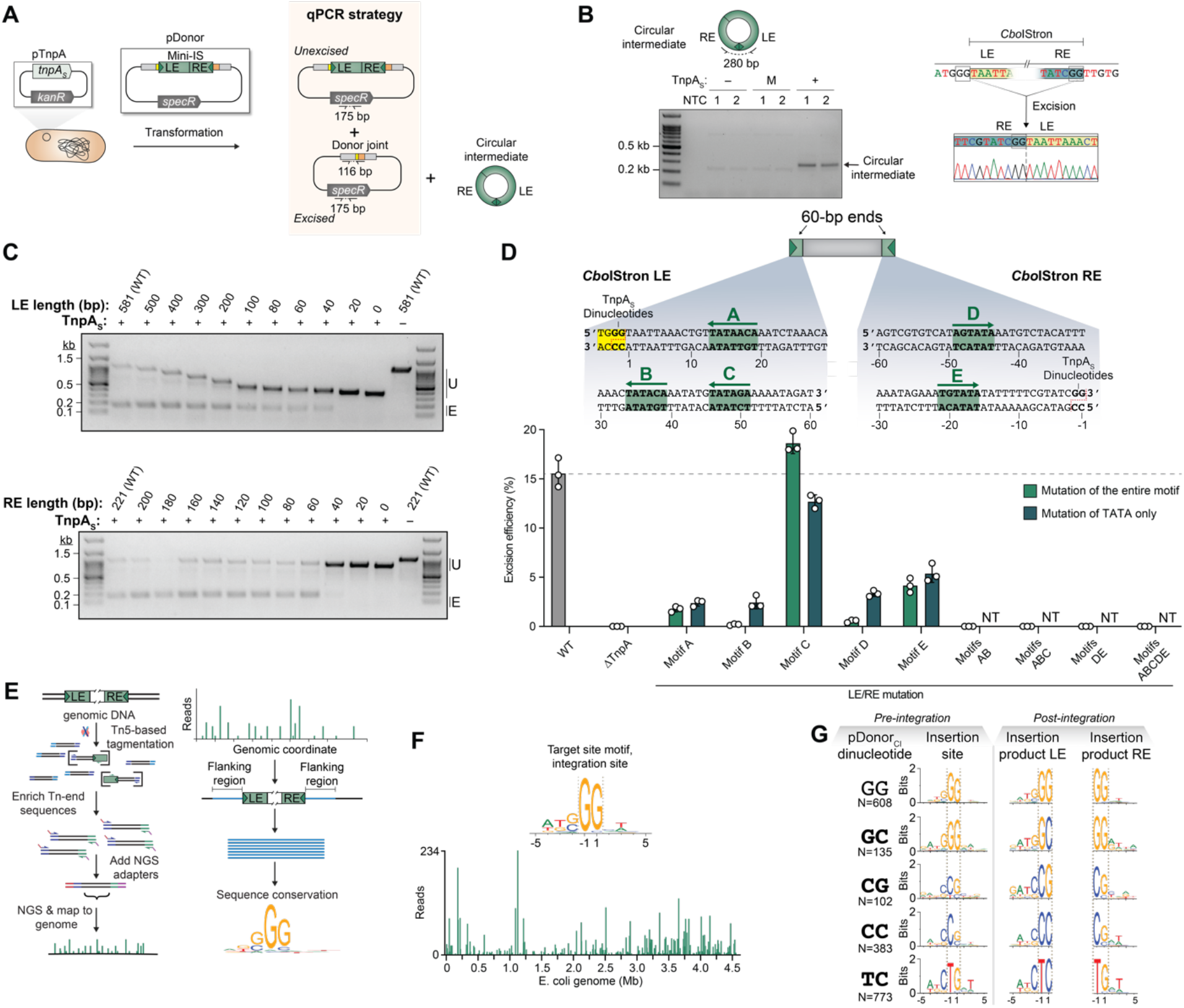
Molecular and sequence determinants of *Cbo*IStron DNA excision and integration target selection by *Cbo*TnpA_S_. (**A**) Schematic of qPCR-based transposon excision assay using a *Cbo*TnpA_S_ expression plasmid (pTnpA_S_) and *Cbo*IStron donor plasmid harboring a mini-transposon with LE and RE boundaries (pDonor). Expected substrates and products generated upon transposon excision are indicated, as are the primer binding sites for quantitative excision measurements using qPCR. (**B)** Gel electrophoresis (left) and Sanger sequencing (right) of PCR products from Fig. 2A, demonstrating the cellular presence of transposon circular intermediates (CircInt) in a TnpA_S_-dependent reaction. Primers were designed to amplify across the joined LE and RE, such that only the indicated product (top) would yield a PCR amplicon 280 bp in size. Reactions were performed in biological duplicates and contained either empty vector (–), mutant S67A TnpA_S_ (M), or WT TnpA_S_ (+); NTC, no-template control. The Sanger sequencing data demonstrate that amplicons contained the inverted RE-LE junction, with recombined GG core dinucleotide. (**C**) Gel electrophoresis of PCR products from experiments performed as in Fig. 2A using mini-IS substrates that contained serial truncations of either the LE (top) or right end (bottom). These experiments indicate that only 40-bp and 60-bp are necessary on the LE and RE, respectively, for WT efficiencies of excision. The length of the truncated end is shown above each lane, counting from the first bp inside the LE or RE, and the mobility of PCR products represent the unexcised (U) or excised (E) products. Note that the excised product is the same size in all cases, as expected. Control lanes on the far right lacked TnpA_S_ and thus were inactive for excision. (**D**) Schematic of minimal transposon design containing 60-bp LE and RE sequences (top); DNA sequence of minimal ends, highlighting the identification of putative TnpA_S_ binding sites denoted A–E (middle); and mini-IS DNA excision assay measured by qPCR (bottom). Binding sites were mutated independently or in tandem, across either the entire motif or only the TATA portion, as indicated. In all cases, disruption of two or more motifs completely abolished detectable DNA integration. Selective TATA mutations were only tested for the indicated constructs; NT, not tested. (**E**) TagTn-seq workflow for deep sequencing of genome-wide transposition events in *E. coli* using TnpA_S_ and circularized intermediate donor molecules (pDonor_CI_). Transposon-containing molecules are selectively amplified in a nested PCR after tagmentation of high-molecular weight genomic DNA, followed by next-generation sequencing (NGS), computational filtering, and read mapping back to the *E. coli* reference genome (left). Meta-analysis of the genomic coordinates containing transposon insertions enables identification of conserved target-site motifs (right). (**F**) Genome-wide distribution of TagTn-seq reads from experiments as in **E**, using WT TnpA_S_ and pDonor_CI_ containing a core GG dinucleotide (colony counts are shown in fig. 2F), mapped to the *E. coli* genome (bottom). Meta-analyses reveal a strict GG dinucleotide requirement at the site of transposon integration (top). (**G**) Experiments in **F** were repeated, but using pDonor_CI_ substrates containing non-canonical core dinucleotides, as indicated. Analysis of the resulting integration sites revealed that integration site preference could only partially be reprogrammed with the altered core dinucleotides; the nucleotide sequence in the LE insertion product always matched the altered core dinucleotide on pDonor_CI_; but a G was preferentially installed at the +1 position in the RE insertion product, regardless of altered core dinucleotide on pDonor_CI_. “N=…” represents the number of unique genomic integration sites detected when sequencing from the left end of the IS element for each core dinucleotide listed.

**Fig. S7.**
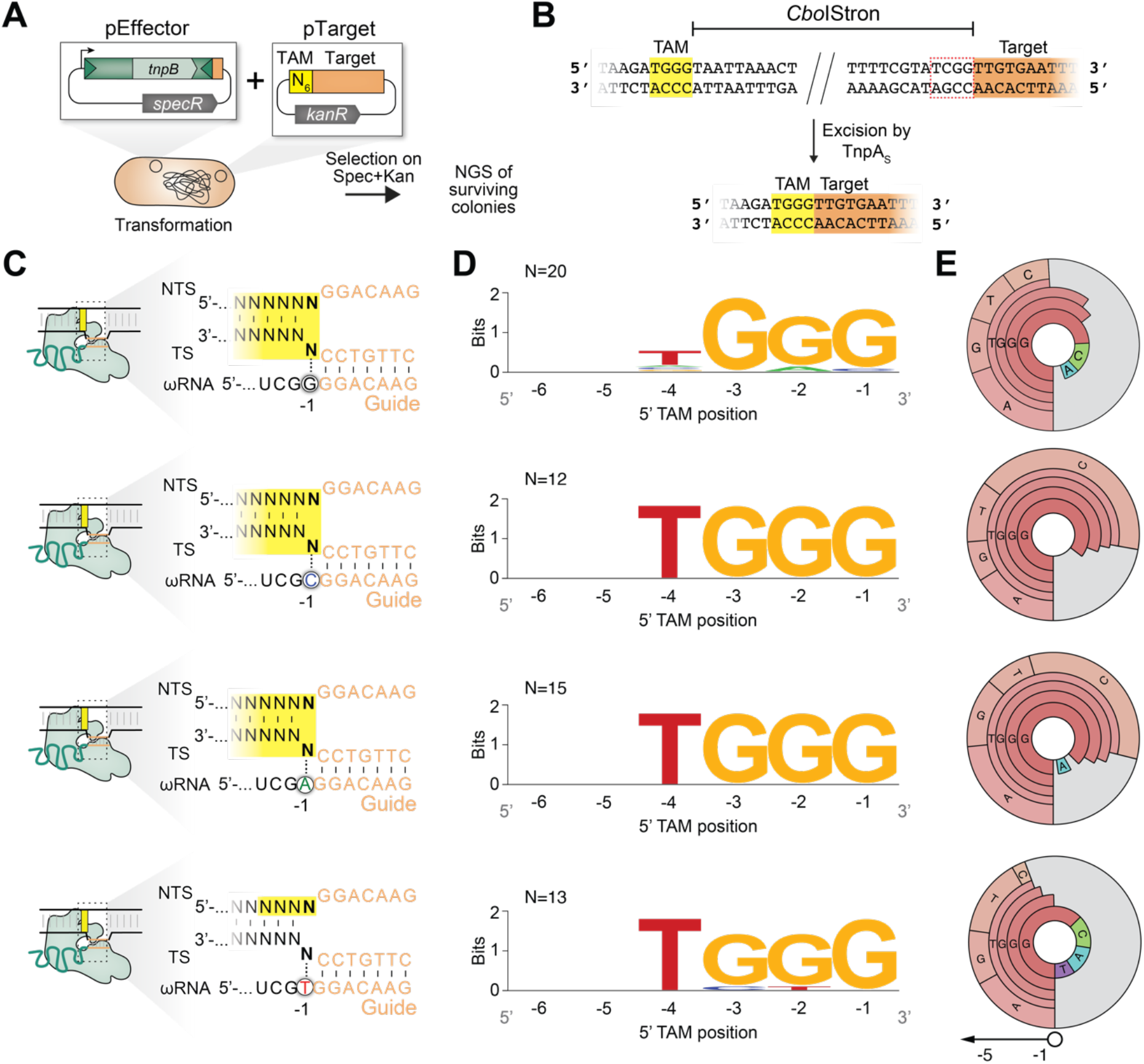
Library experiments to determine TAM specificity by *Cbo*TnpB. (**A**) Schematic of TAM library cleavage assay. A plasmid expressing nuclease-active *Cbo*TnpB and an associated ωRNA from within the native *Cbo*IStron (pEffector) is designed to cleave a target sequence adjacent to a randomized 6-mer (pTarget). Plasmid cleavage results in plasmid elimination, loss of cell viability during drug selection, and depletion of the particular TAM upon library sequencing. (**B**) Schematic highlighting the similarities between the LE-adjacent TAM (5′-TGGG-3′, yellow) and the target-adjacent sequence within the transposon RE (5′-TCGG-3′, red dotted line), from which the ωRNA is transcribed. ‘Self’ versus ‘non-self’ discrimination between the ωRNA template and post-excision donor joint, respectively, requires stringent discrimination at the –3 position. (**C**) WT (top) and non-canonical ωRNA variants screened in the TAM library assay, to investigate if base-pairing occurs at the –1 position in the ωRNA. NTS, non-target strand; TS, target strand. Randomized TAM sequence is highlighted in yellow. (**D**) WebLogo representation of top depleted library members for the ωRNA variants shown in panel **B**; The number of library members used to construct the WebLogo is shown in the top left corner. Data for the WT ωRNA are replotted from Fig. 3F. (**E**) TAM wheels for the same ωRNA variants shown in **B**, generated using the 5% most depleted library members. These results indicate that the –1 position of the ωRNA does not confer any specificity for the recognition of TAM motif.

**Fig. S8.**
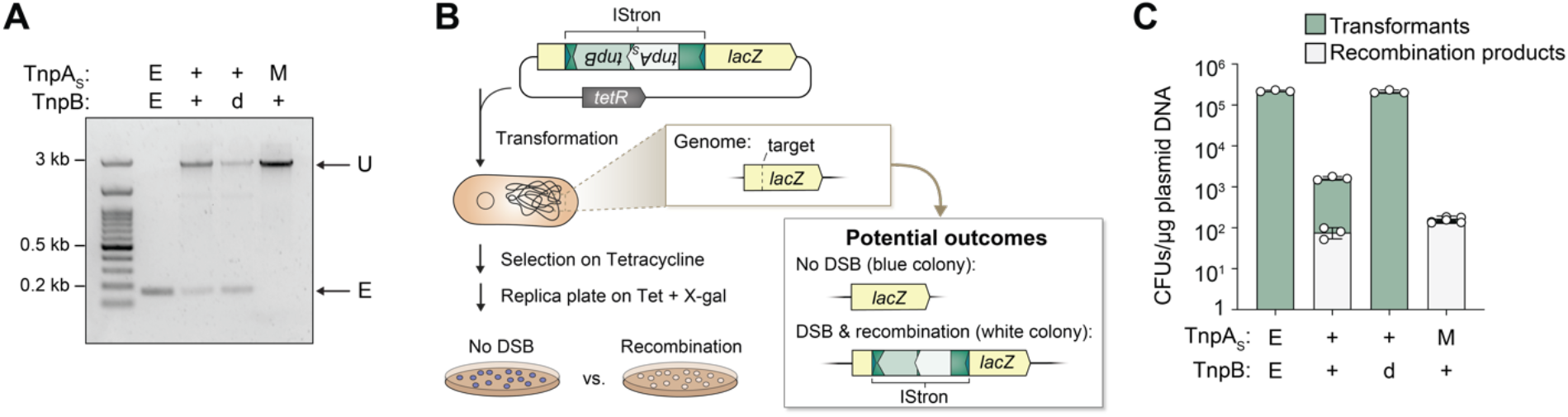
Synergistic TnpA_S_–TnpB activity during transposon integration and recombination. (**A**) PCR and gel analysis of *lacZ* genotypes from Fig. 3I, demonstrating the role of TnpB in promoting transposon retention by reducing the relative frequency of excision products (E) relative to unexcised transposon substrates (U). Cells expressed either wild-type (+) or mutant (S67A) TnpA (M), in the presence of WT (+) or nuclease-dead (D189A) TnpB (d). (**B**) Workflow to measure transposon recombination in *E. coli* with TnpA_S_ and TnpB. Native *Cbo*IStron transposons with TnpA_S_ and either WT or nuclease-dead TnpB were inserted in the reverse direction at a compatible TAM in plasmid-encoded *lacZ*, such that splicing could not generate a *lacZ*+ phenotype. Plasmids were used to transform *E. coli* cells harboring a wild-type *lacZ* locus. RNA-guided DNA cleavage of genomic *lacZ* triggers recombination with the ectopic *Cbo*IStron-*lacZ*, leading to white colonies. Tet, tetracycline. (**C**) Bar graph shows the plasmid transformation efficiency for each condition, with white bars reporting white colony phenotype; E, empty vector. The data reveal that TnpA_S_ and TnpB co-operate for efficient self-mobilization into a vacant donor site via recombination, but only in the presence of nuclease-active TnpB. Data are mean ± s.d. (n = 3).

**Fig. S9.**
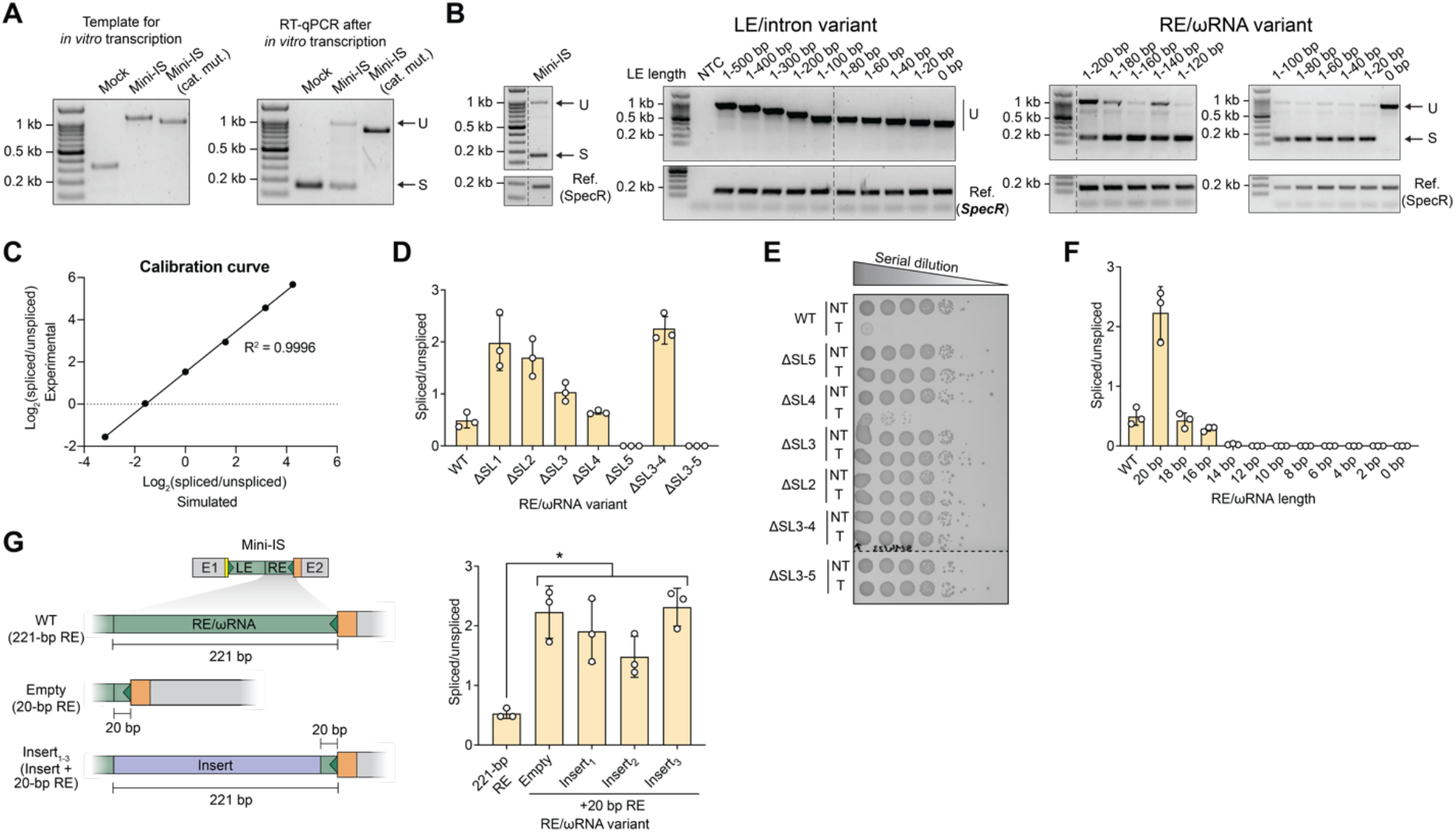
Splicing and RNA-guided DNA cleavage activity for modified IStron variants. (**A**) Templates for in vitro transcription (IVT)-based group I intron splicing assays were generated by PCR and lacked any detectable truncation products (left). The ensuing IVT reactions revealed evidence of spliced exon-exon junction products, as detected by RT-qPCR (right), which matched the expected size based on a mock control. IStron (cat. mut.) contains a P7-P9 loop deletion in the intron catalytic core; U, unspliced; S, spliced. (**B**) Agarose gel electrophoresis of RT-PCR products from splicing assays with the indicated serial deletions in the transposon left end/intron region (LE/intron, left) or transposon right end/ωRNA region (RE/ωRNA, right). Unspliced (U) and spliced (S) products are indicated, relative to reference amplicons for a SpecR drug marker (bottom). Any deletion in the 581-bp LE/intron region eliminates splicing, whereas deletions of everything but the terminal 20 bp in the RE/ωRNA region are tolerated. NTC, non-template control. (**C**) Comparison of simulated spliced/unspliced ratios, generated by mixing mock-spliced and mock-unspliced lysates in known ratios, versus experimentally determined spliced-unspliced ratios measured by RT-qPCR, using the strategy described in Fig. 4A. The results demonstrate the accuracy of our quantification method. (**D**) Quantitative measurements of the spliced/unspliced ratio by RT-qPCR for the indicated constructs that harbor stem-loop (SL) deletions in the RE/ωRNA region, as defined in Fig. 5A. The WT construct contains 221-bp of the RE. (**E**) Bacterial spot assays for the same RE/ωRNA SL deletion constructs in **D**, in which RNA-guided DNA cleavage leads to cell death. Cells expressing TnpB were transformed with either target-bearing (T) or no target-bearing (NT) plasmid, and transformants were serially diluted, plated on selective media, and cultured at 37 °C for 24 hours. Deletion of any SL (except for SL4) completely abolished DNA cleavage activity. An SL1 deletion was not tested due to its overlap with the *tnpB* ORF. (**F**) Quantitative measurements of the spliced/unspliced ratio by RT-qPCR for the indicated constructs that harbor focused deletions in the 3′-terminal RE/ωRNA region. The WT construct contains 221-bp of the RE, however a shorter 20-bp construct was shown to exhibit far greater splicing activity. Any deletion beyond 16 bp leads to a loss of splicing activity. (**G**) Schematic (left) and quantitative measurements of the spliced/unspliced ratio by RT-qPCR (right) for the indicated constructs that harbor one of three *lacZ*-derived insertions (Insert_1–3_). Removal of the ωRNA altogether for the 20-bp RE variant (Empty), or its replacement with these insertions, all stimulated splicing efficiency to a similar extent, relative to the 221-bp full-length RE sequence. Statistical significance was determined using Welch’s t-test, * denotes p-value <0.05.

**Fig. S10.**
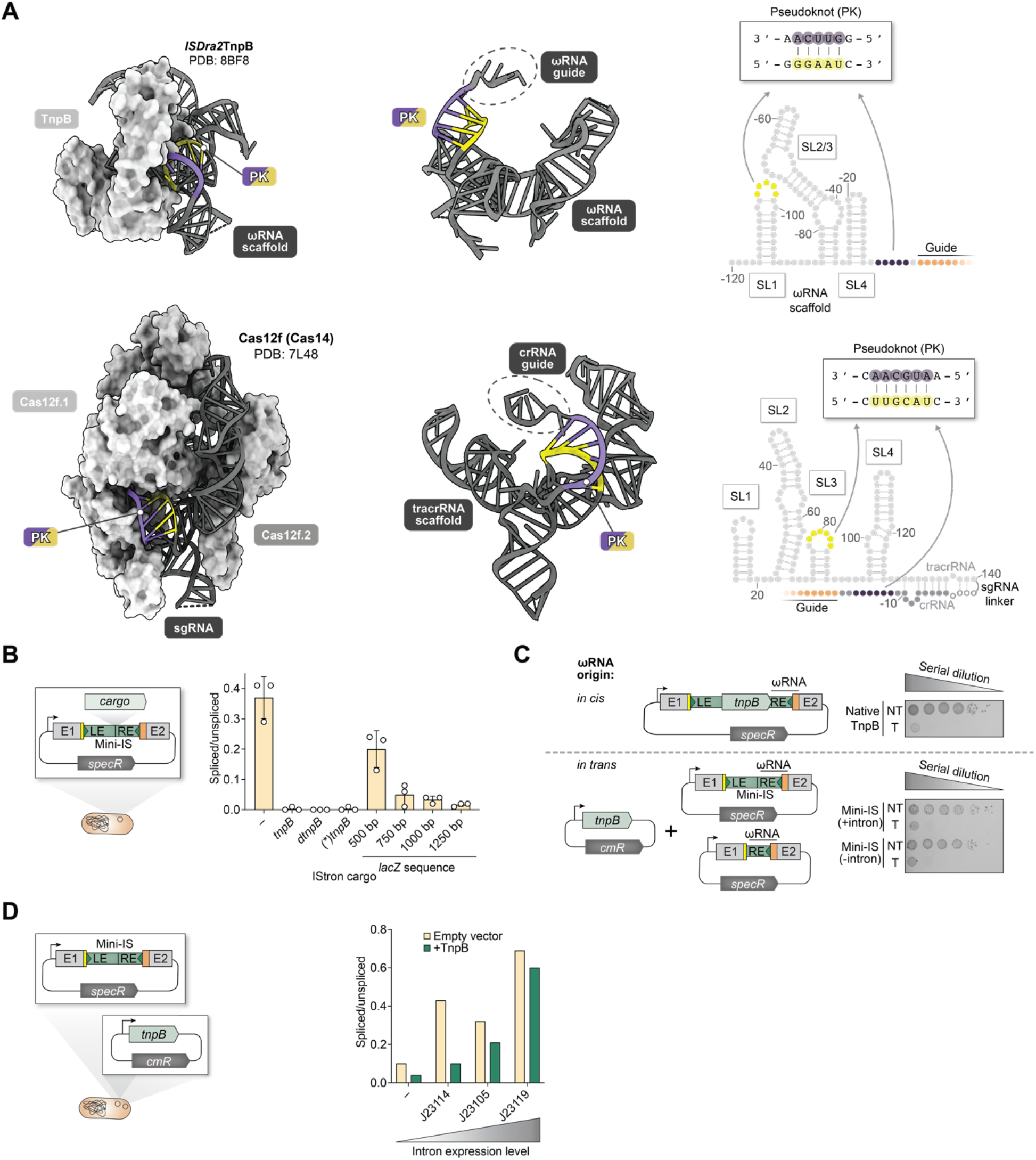
Effect of ωRNA perturbations on the IStron splicing efficiency. (**A**) CryoEM structures of DraTnpB (PDB ID: 8BF8) (*58*) and Cas12f (PDB ID: 7L48) (*99*) nucleases bound to ωRNA and sgRNA, respectively (left), and the same respective RNA molecules shown without the protein (middle). Secondary structures of each guide RNA are shown on the right. The key pseudoknot region is colored in yellow and purple, and the guide sequence is shown in orange. (**B**) Quantitative measurements of the spliced/unspliced ratio by RT-qPCR for the indicated constructs that contain variable ‘cargo’ sequences. The minimal construct harbors the *Cbo*IStron sequence after removal of *tnpA_S_* and *tnpB* ORFs and exhibits a splicing ratio of ∼0.4, whereas splicing becomes nearly undetectable with cargos comprising either the *tnpB* gene (encoding functional TnpB), *dtnpB*, or a *tnpB* with in-frame stop codon ((*)*dtnpB*), all of which are ∼1180 bp in length. Constructs with alternative cargos containing the indicated length of unrelated *lacZ* sequence also exhibited decreased splicing efficiency with increasing size. (**C**) Bacterial spot assays demonstrate that TnpB is equally active for RNA-guided DNA cleavage when the ωRNA is expressed *in cis* with TnpB from a native *Cbo*IStron context (top), or *in trans* from a separate ωRNA expression plasmid (bottom). The *in-trans* activity was equivalent regardless if the mini-IS also encoded the full-length group I intron. Transformants were serially diluted, plated on selective media, and cultured at 37 °C for 24 hours. Data for the *in cis* configuration are copied from Fig. 3D. (**D**) Plasmid design for measuring splicing in the presence of a TnpB expression plasmid *in trans* (left), and bar graph with quantitative measurements of the spliced/unspliced ratio by RT-qPCR for the indicated intron expression levels (right). Variable-strength promoters were used, with (green) or without (yellow) TnpB co-expression. The repressive effect of TnpB is strongest at low expression levels. “–” refers to a construct lacking a synthetic promoter upstream of the intron.

**Fig. S11.**
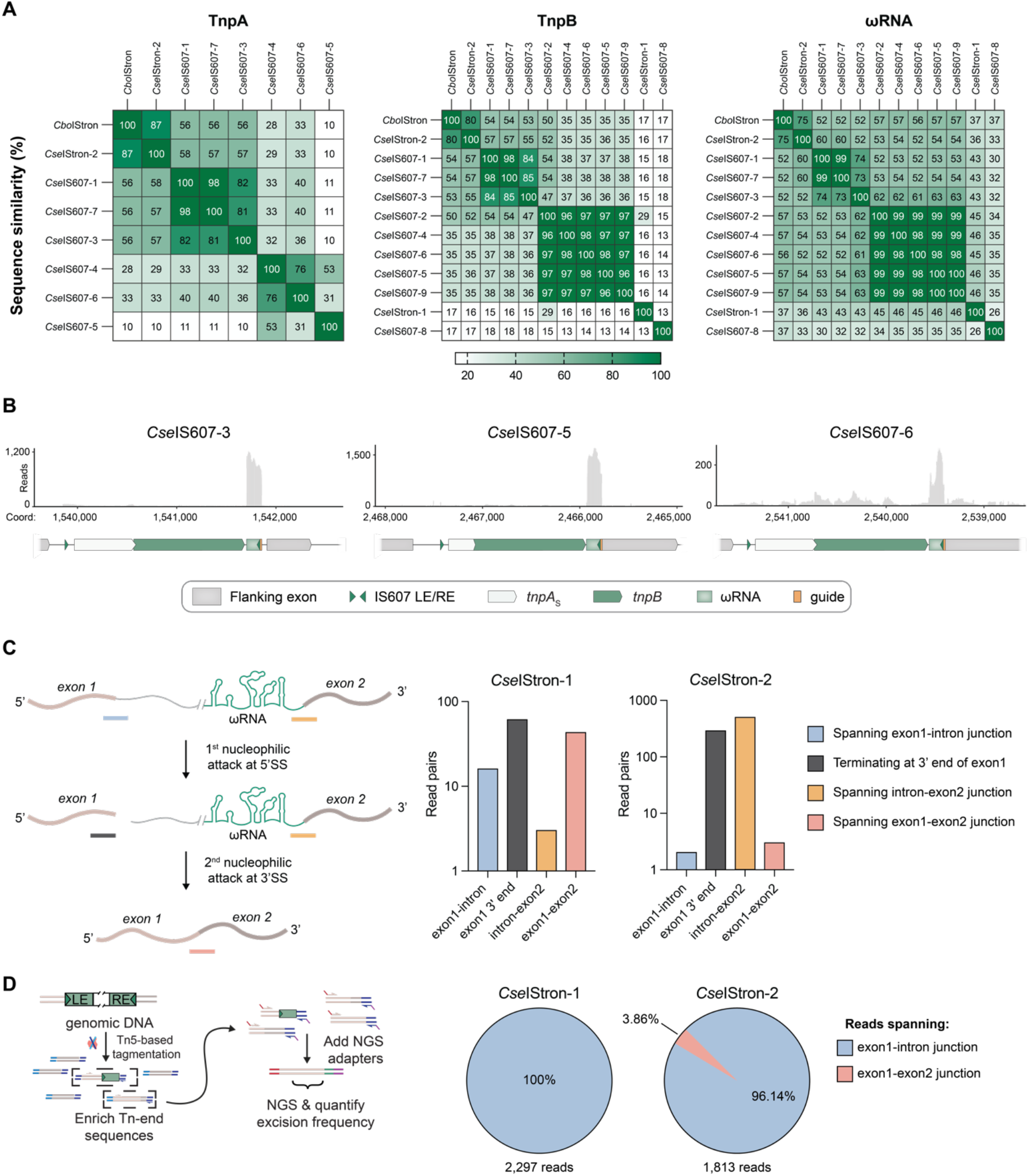
Comparative sequence and RNA-seq analyses of *C. senegalense* IS607 and IStron elements. (**A**) Pairwise sequence identity matrices for all identifiable TnpA (left) and TnpB homologs (middle), and ωRNAs (right), encoded by *C. senegalense* strain (ATCC 25772), as well as experimentally tested *Cbo*IStron. Percent identities are shown in the heat maps, with row and column orders determined by hierarchical clustering. TnpA and TnpB were compared using protein amino acid sequence, ωRNAs – using nucleotide sequence. (**B**) Total RNA-seq read coverage over representative *Cse*IS607 (intron-less) loci. The genomic coordinates are shown, as are gene annotations below each graph. (**C**) A schematic of the two-step transesterification reaction underlying group I intron splicing is shown (left). Mapping of read pairs to unspliced, partially spliced, or fully spliced transcripts is quantitated for each *Cse*IStron locus (right). The actual read pair count for the two *Cse*IStrons (right) reveals that while *Cse*IStron-1 is splicing efficiently, *Cse*IStron-2 is likely undergoing incomplete splicing. (**D**) Targeted genomic IStron sequencing workflow (left) and proportion of exon1-intron junction reads versus exon1-exon2 junction spanning reads for both *Cse*IStrons (right). Presence of exon1-exon2 reads in genomic DNA for *Cse*IStron-2 loci indicates that the element is excised from some copies of the genome.

## SUPPLEMENTARY TABLES

**Table S1.** Group I introns, IS605- and IS607-family elements from Clostridium species used in this study.

**Table S2.** Strains used in this study.

**Table S3:** Description and sequence of plasmids used in this study.

**Table S4:** Oligonucleotides used in this study.

